# Deciphering the Antigenic Evolution of Seasonal Influenza A Viruses with PREDAC-Transformer: From Antigenic Clustering to Key Site Identification

**DOI:** 10.1101/2025.10.29.684974

**Authors:** Jingze Liu, Jiangyuan Wang, Chuan Wang, Xiao Ding, Wenping Xie, Luyao Qin, Aiping Wu, Jing Meng, Taijiao Jiang

**Author notes:** These authors contributed equally: Jingze Liu, Jiangyuan Wang, Chuan Wang.

## Abstract

Seasonal influenza viruses undergo continuous antigenic drift due to mutations in the hemagglutinin (HA) protein, rendering vaccines ineffective and posing a significant global public health challenge. Existing computational models can predict antigenic relationships but generally lack interpretability, making it difficult to reveal the molecular determinants driving antigenic changes. To address this, we developed PREDAC-Transformer, an end-to-end deep learning framework that integrates sequence, physicochemical, and evolutionary features, and employs a self-attention mechanism to capture long-range dependencies relevant to antigenicity. More importantly, we introduce the integrated antigenicity score, which combines attention-based attribution with information-theoretic metrics to provide continuous quantification and ranking of the antigenic contributions of individual amino acid site. Our results demonstrate that PREDAC-Transformer not only significantly improves the accuracy of antigenic relationship prediction but also successfully recapitulates major historical antigenic cluster transitions. Using integrated antigenicity score, we systematically identified two classes of key sites: global key sites with sustained impact on antigenic evolution, and cluster-transition determining sites that drive cluster transitions. These sites include most canonical epitopes and reveal additional functional residues previously overlooked, which may influence immune escape via cooperative effects or glycosylation. Collectively, these findings advance our understanding of influenza antigenic evolution and provide novel insights for refining computational models. PREDAC-Transformer achieves high-precision prediction while attributing antigenic differences to individual residues, thereby linking viral genomic variation, antigenic change, and public health decision-making. This framework has the potential to reduce experimental burdens in influenza surveillance and assist in vaccine strain recommendation, thereby supporting global influenza control efforts.

## Introduction

Seasonal influenza A viruses are responsible for millions of infections and hundreds of thousands of deaths annually, posing a persistent threat to global public health (Gilbertson and Subbarao 2023). Vaccination remains the most effective strategy for influenza prevention and control. Each year, the World Health Organization (WHO) recommends vaccine strains for the upcoming season based on surveillance data. However, the efficacy of these vaccines is contingent on the antigenic congruence between the vaccine strain and circulating viruses (Trombetta et al. 2022). The surface hemagglutinin (HA) protein of the influenza virus is the main target of neutralizing antibodies, which undergoes antigenic drift through the continuous accumulation of amino acid mutations, thereby compromising vaccine effectiveness (Clark et al. 2024). Therefore, the accurate prediction of antigenic relationship and a thorough understanding of its molecular underpinnings are pivotal for optimizing vaccine strain recommendation.

The traditional hemagglutination inhibition (HI) assay is the standard for measuring antigenicity, yet it is time-consuming, labor-intensive, and can yield inconsistent results across laboratories (Zacour et al. 2016; Laszlofy et al. 2024). Smith et al. used multidimensional scaling (MDS) of HI data to demonstrate that the antigenic evolution of influenza A/H3N2 virus occurs in a distinct, cluster-wise manner, identifying 11 such clusters between 1968 and 2003 (Smith et al. 2004). This work spurred the development of computational methods to predict the antigenicity of emerging strains and infer antigenic clusters. Early computational models primarily employed machine learning approaches to predict antigenic relationships. For example, the studies by Liao (Liao et al. 2008), Lee (Lee and Chen 2004), and Du (Du et al. 2012) focused on antigenic epitopes and the physicochemical properties of amino acids to score or predict antigenic relationships. However, these approaches were limited as they only considered a small number of amino acid sites and could not fully assess the collective impact of point mutations across the entire HA1 sequence on antigenicity. More recently, deep learning methods have been applied to antigenic relationship prediction of influenza virus, owing to their capacity to learn complex features and model interactions between amino acid sites. Following the initial application of Convolutional Neural Networks (CNNs) by Lee et al. (Lee et al. 2020), several other models were developed, including FLU (Forghani and Khachay 2020), IAV-CNN (Yin et al. 2022), and our own PREDAC-CNN (Meng et al. 2024) and PREDAC-FluB (Xie et al. 2025). While these CNN-based models excel at capturing local dependencies, their convolutional architecture is less effective at integrating long-range interactions, which are crucial for characterizing the synergistic effects of sites that form conformational epitopes. Furthermore, despite their predictive accuracy, these models operate as black boxes. They are able to predict antigenic relationships but cannot pinpoint the specific amino acid substitutions driving these differences, and therefore lack interpretability.

To elucidate the molecular mechanisms underlying antigenic variation, subsequent research has focused on identifying key amino acid sites that drive antigenic evolution. Experimental studies, such as those by Koel et al. (Koel et al. 2013), have pinpointed a few critical residues responsible for major antigenic cluster transitions in H3N2 viruses, but these experiments are costly and difficult to apply systematically across all viral lineages. To overcome these limitations, computational approaches have been developed to infer sites associated with antigenic change—for example, the works of Steinrueck and McHardy (Steinbrück and McHardy 2012) and Neher et al. (Neher et al. 2016). Our previous method, RECDS (Quan et al. 2019), used traditional machine learning model contributions to identify cluster-transition determining sites, but global averaging of scores often diluted the signal, capturing only a few sites while overlooking many with moderate effects or synergistic interactions. Similarly, approaches based on information gain or Shannon entropy (Cheng et al. 2024; Zhai et al. 2024), such as those by Cheng et al. (Cheng et al. 2024), rely on indirect, model-dependent importance metrics. In all these cases, the importance scores quantify a site’s contribution to the model’s classification decisions rather than directly capturing its impact on the magnitude of antigenic change. Consequently, the precise molecular determinants of antigenic differences between strains remain unresolved, unlike in Transformer-based models where attention weights can be learned end-to-end from the sequence data (Meng et al. 2025).

To address these gaps in prediction and interpretation, we developed PREDAC-Transformer, a novel analytical framework. The framework’s core advantages are reflected in three aspects: First, it employs a Transformer encoder (Vaswani et al. 2017), leveraging self-attention mechanisms to effectively capture the long-range dependencies that determine antigenicity, overcoming the local-feature limitations of CNNs. Second, it utilizes a composite feature representation that combines raw sequence data, physicochemical properties, and ESM-2 embeddings from a large-scale protein language model (Lin et al. 2023), allowing it to integrate sequential, chemical, and evolutionary information. Third, and most critically, we propose the integrated antigenicity score, derived from the model’s attention weights, to identify key amino acid sites driving antigenic variation. The integrated antigenicity score combines model-based attribution with data-driven statistical significance, effectively reducing false positives that arise from model artifact or mutations at sites irrelevant to antigenicity.

Our results show that PREDAC-Transformer significantly improves the accuracy of antigenic prediction and successfully recapitulates major historical antigenic shift events for both H3N2 and H1N1 viruses. Through the integrated antigenicity score, we systematically identified two categories of functional sites: global key sites that play a sustained role in long-term evolution, and cluster-transition determining sites that are decisive during the transition between antigenic clusters. In addition to validating known critical sites, we discovered novel sites outside of traditional epitopes that significantly impact antigenicity, potentially through synergistic interactions or mechanisms such as glycosylation masking. By connecting high-precision prediction with precise attribution of antigenic contributions to individual amino acid sites, this framework creates a seamless link between genomic variation, antigenic evolution, and public health decision-making. PREDAC-Transformer holds the potential to reduce the burden of experimental surveillance and enhance the accuracy of vaccine strain recommendations, providing robust support for global influenza prevention and control.

## Materials and Methods

### Datasets and Preprocessing

In this study, HI assay data were gathered from the WHO and various national health agencies. Corresponding HA sequences were sourced from the Global Initiative on Sharing All Influenza Data (GISAID) (Shu and McCauley 2017) and the NCBI Influenza Virus Database (Bao et al. 2008). To ensure data quality, we first excluded records with missing or ambiguous HI titer values. We then removed virus-antiserum titration records involving the same viral strain, as these do not inform antigenic differences. Titration data with conflicting antigenicity results were also discarded to maintain consistency. The final dataset for model training comprised 8,962 strain pairs for influenza A/H3N2 viruses (1968–2022) and 11,038 strain pairs for A/H1N1 viruses (1995–2024).

For the inference of antigenic clusters, we obtained HA1 sequences of influenza A/H3N2 (1968– 2024) and A/H1N1 (1977–2024) viruses from GISAID. The preprocessing pipeline involved several quality control steps: sequences with lengths deviating more than 10% from the canonical length or containing over 1% ambiguous amino acids were removed. We retained only strains isolated from human hosts with clearly documented collection years. After this initial filtering, 120,596 A/H3N2 and 98,608 A/H1N1 sequences remained. We then deduplicated sequences based on their earliest year of collection. Subsequently, the sequences were deduplicated. CD-HIT (Li and Godzik 2006) was employed to reduce sequence redundancy annually (similarity threshold: 0.988), resulting in 3,289 representative HA1 sequences for A/H3N2 and 3,051 for A/H1N1. Finally, a phylogenetic tree was constructed using FastTree (Price et al. 2009), and any outlier sequences were removed, yielding a final count of 3,283 H3N2 and 3,037 H1N1 sequences. These procedures ensured a high-quality and representative sequence dataset for the subsequent phylogenetic analysis and inference of antigenic clusters.

### Characterization of antigenic relationships of strain pairs

Antigenic distance was calculated using both bidirectional and unidirectional methods. The bidirectional distance was determined using the Archetti-Horsfall formula (Ndifon et al. 2009):

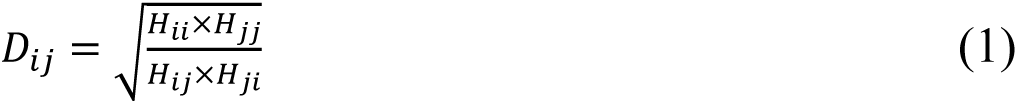

In this context, *𝐻_ij_* is the HI titer of strain i needed to inhibit cell agglutination caused by strain j. A distance value greater than or equal to 4 indicates that two strains are antigenic variants, while a value less than 4 suggests they are antigenically similar. In cases where homologous titers were unavailable for a strain pair (for example, only 900 bidirectional pairs are available for H1N1), a unidirectional antigenic distance was calculated as the ratio of the heterologous HI titer to the homologous titer *𝐷_ij_ = 𝐻_jj_/𝐻_ij_* (Neher et al. 2016).

Based on these criteria, the 8,962 strain pairs for A/H3N2 viruses were categorized into 4,966 antigenically distinct (positive) pairs and 3,996 antigenically similar (negative) pairs. For the 11,038 A/H1N1 strain pairs, 3,633 were classified as antigenically distinct (positive) and 7,405 as antigenically similar (negative). Supplementary Tables S1 and S2 detail the yearly distribution of these strain pairs. In these analyses, each strain pair consists of one strain isolated in a specific year and another strain isolated in or before that year.

### Workflow of PREDAC-Transformer

The PREDAC-Transformer workflow is illustrated in Figure 1. The model takes paired HA1 sequences of influenza A/H3N2 or A/H1N1 viruses as input. We first designed a hybrid input representation that combines physicochemical properties of amino acids with contextual embeddings generated by the ESM-2 protein language model. These features for the paired HA1 sequences are integrated into an input matrix tailored for the Transformer architecture. Utilizing this input matrix, a Transformer encoder-based model predicts the antigenic relationship between the paired strains. The model’s final Softmax probabilities are then used to construct an antigenic distance matrix. Subsequently, we apply UMAP for dimensionality reduction followed by K-means clustering to infer antigenic clusters (Xie et al. 2025).

**Figure 1.**
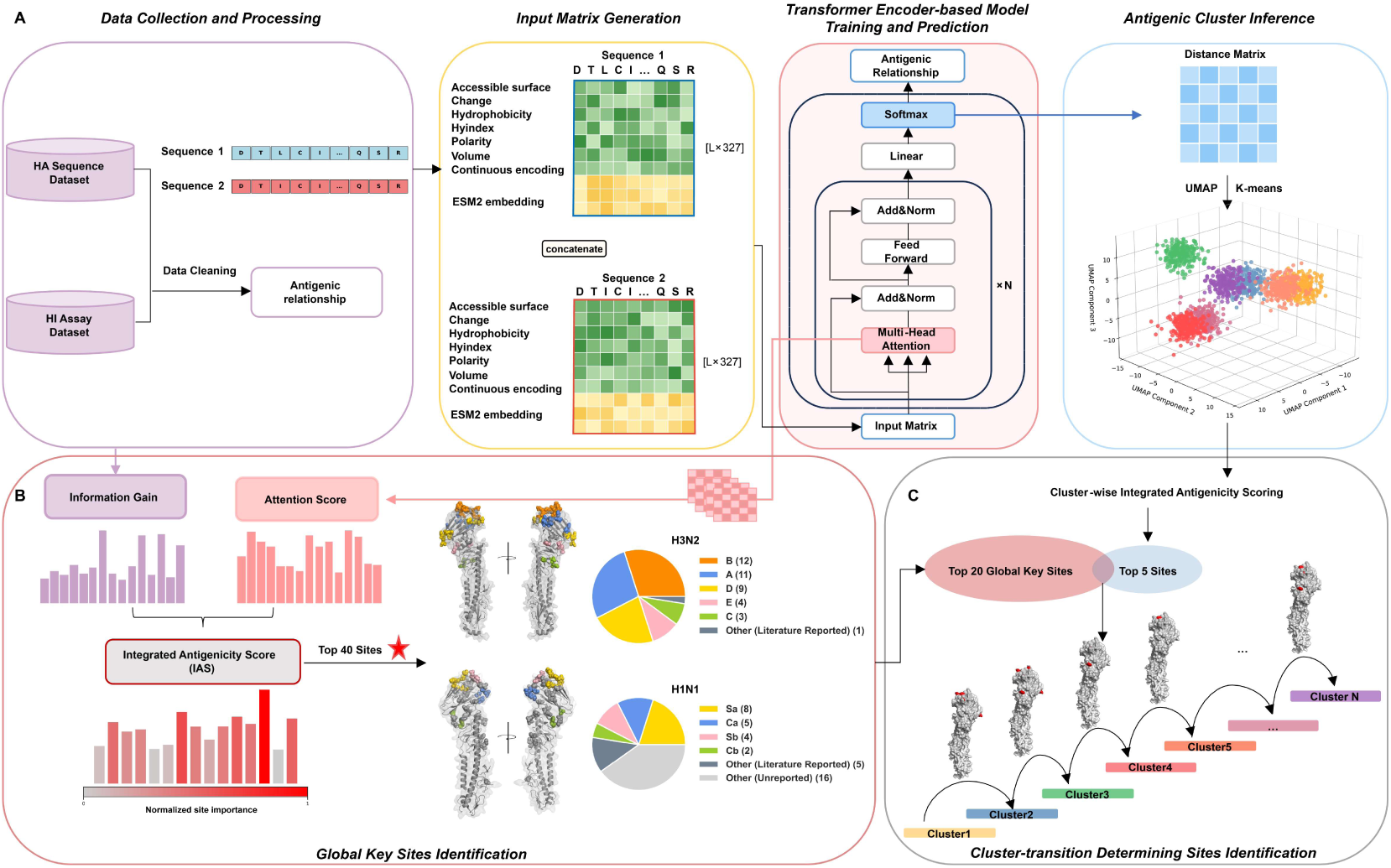
The PREDAC-Transformer framework and its application in identifying key antigenic sites. (A) Overview of the PREDAC-Transformer workflow. PREDAC-Transformer analyzes paired HA1 sequences from influenza A/H3N2 or A/H1N1 viruses. Sequences are encoded with selected physicochemical properties and contextual embeddings from ESM-2, processed by the Transformer encoder to predict antigenic relationships. Prediction probabilities are used to construct a distance matrix, and antigenic clusters are identified via K-means after UMAP dimensionality reduction. (B) Workflow for identifying globally important antigenic sites. Information gain and model-derived attention scores are combined to calculate an integrated antigenicity score for each amino acid position. The 40 sites with the highest integrated antigenicity score are visualized on the three-dimensional HA structures of H3N2 (PDB: 2YPG) and H1N1 (PDB: 2WRG) viruses. The accompanying pie charts quantify the overlap between these model-identified sites and canonical antigenic sites (A–E for H3N2; Sa, Sb, Ca, Cb for H1N1) from existing literature. (C) Methodology for identifying cluster-transition determining sites. Antigenic clusters are first identified from a large HA sequence dataset using the model’s predictions (as described in panel A). The integrated antigenicity score is then specifically calculated for strain pairs spanning adjacent clusters to pinpoint amino acid substitutions critical for the evolutionary shift, illustrating the molecular pathway of antigenic evolution.

#### Hybrid Input Representation

To comprehensively characterize the HA1 sequence, we developed a hybrid input representation method. This approach integrates two types of features. First, we adopted the seven features used in PREDAC-CNN (Meng et al. 2024): six key physicochemical properties of amino acids (accessible surface, charge, hydrophobicity, hydrogen bond index, polarity, and volume) and a continuous encoding for the 20 standard amino acids. Second, we incorporated contextual embeddings for each amino acid generated by the large-scale protein language model ESM-2 (8M). Pre-trained on large-scale protein sequence data, ESM-2 embeddings capture both the intrinsic properties of individual amino acids and their contextual relationships within sequences, thereby providing biologically meaningful features (Lin et al. 2023). Specifically, for each amino acid, we concatenated the 7-dimensional physicochemical feature vector with the 320-dimensional ESM-2 embedding to create a 327-dimensional input vector. For a pair of HA1 sequences, these vectors are combined to form the final input matrix. The resulting matrix has dimensions of 329×654 for A/H3N2 viruses (329 amino acid sites, 654 feature dimensions per site) and 327×654 for A/H1N1 viruses. This hybrid approach retains the interpretable physicochemical features while leveraging the rich contextual information from the ESM-2 model.

#### The Transformer Encoder-Based Model for Prediction

The high dimensionality of the input representation presents challenges for traditional machine learning algorithms. While Convolutional Neural Networks (CNNs) are adept at capturing local features, they lack interpretability and are less effective at modeling long-range dependencies within sequences. To address this limitation, we adopted a Transformer encoder architecture to comprehensively model global dependencies across the HA1 sequence. The Transformer encoder is composed of stacked multi-head self-attention and feed-forward network layers, with residual connections and layer normalization to ensure stable training (Vaswani et al. 2017). The multi-head self-attention mechanism allows the model to simultaneously learn dependencies between all pairs of positions in the sequence, effectively balancing the contributions of local mutations and global cooperative effects. Furthermore, the attention weights offer a degree of interpretability by indicating the relative importance of each amino acid site in the prediction.

We utilize this Transformer encoder to ascertain the antigenic relationship between two strains of either influenza A/H3N2 or A/H1N1 viruses. The detailed model architecture is shown in Figure 1. We used a two-layer Transformer encoder with an embedding dimension of 16 and a feed-forward network dimension of 64. The model for A/H3N2 employed 8 attention heads, while the model for A/H1N1 used 4. The model was trained using the AdamW optimizer (learning rate = 1e-3, weight decay = 1e-8) with an early stopping criterion. The detailed comparison of model hyperparameters can be found in Supplementary Figure S1. The output from the Transformer encoder is passed to a classification head consisting of fully connected layers, and a final Softmax layer computes the probability of antigenic variation.

#### Antigenic cluster inference

To infer antigenic clusters, we first constructed a pairwise antigenic distance matrix using the antigenic variation probabilities from the model’s Softmax output. Since the volume of data required for clustering is large, we assumed that if the year difference between two strains exceeded 15 years, they would be considered antigenically distinct, with a probability of 1. We then applied Uniform Manifold Approximation and Projection (UMAP) to reduce the dimensionality of this distance matrix to three, preserving the underlying cluster structure while reducing complexity (McInnes et al. 2020; Xie et al. 2025). Finally, K-means clustering was performed to partition the data into antigenic clusters. The optimal number of clusters (k) was determined using the elbow method, which identifies the optimal clustering number by analyzing the relationship between the within-cluster sum of squares (WCSS) and the number of clusters (Thorndike 1953; Syakur et al. 2018). This analysis suggested an initial value of k=28 for A/H3N2 and k=7 for A/H1N1. A secondary clustering was performed on the sparse, early A/H1N1 data (pre-2009), for which the elbow method indicated an optimal k=11. This approach enabled the identification of major antigenic clusters for A/H3N2 (1968–2024) and A/H1N1 (1977–2024) viruses (Supplementary Figure S6). Circulation time is defined as clusters appearing in three or more consecutive years, excluding any gaps of one or more years.

### Identification of Key Antigenic Sites

#### Derivation of the Integrated Antigenicity Score

To identify the key amino acid sites driving antigenic variation, we developed a multi-level analysis strategy based on a novel metric, the integrated antigenicity score. The integrated antigenicity score combines two key components: model attribution, derived from Transformer attention weights, and data-driven statistical significance, quantified by information gain. Information gain is used to identify key features by quantifying the contribution of each site in reducing data uncertainty, thereby identifying the most discriminative sites for a given classification task (Quinlan 1986; Alhaj et al. 2016; Cheng et al. 2024). The calculation of this information gain is task-specific: it is derived from the antigenic relationships in HI assay data across the evolutionary history for global analysis or from antigenic cluster membership for cluster-transition analysis. This dual-component design ensures that only sites exhibiting strong signals in both model interpretation and statistical analysis receive high scores, thereby effectively filtering out two common sources of noise: false-positive sites arising from attribution bias with low information gain, and sites with frequent mutations but limited antigenic impact, characterized by low attention. The integrated antigenicity score is defined as:

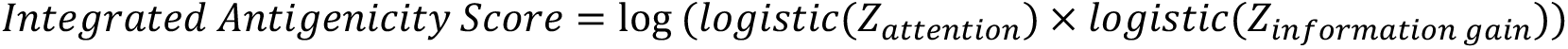

Here, *𝑍_attention_* is the z-score normalized attention weight, aggregated across all encoder layers and attention heads, which quantifies the model’s focus on a given site (Vaswani et al. 2017). *𝑍_information gain_* is the z-score normalized information gain, measuring the discriminatory power of an amino acid substitution at a site with respect to the classification labels of the task, which may correspond to either antigenic relationship or cluster membership across the evolutionary history. The logistic function was incorporated following the multidimensional scoring strategy employed in the EVEscape model (Thadani et al. 2023), assuming equal weights for the two components. This design ensures dual consistency, such that only sites exhibiting both strong model attention and statistical significance are assigned elevated integrated antigenicity score. It is important to note that the two components of integrated antigenicity score are context-dependent, and their computation may vary depending on the research task. In this study, we operationalized them in two distinct scenarios: the identification of global key sites with sustained impact on antigenic evolution, and the identification of cluster-transition determining sites, the overall workflow is depicted in Figures 1B and 1C.

#### Identification of global key sites

The identification of global key sites aimed to capture residues exerting long-term and widespread influence on antigenic evolution. In this task, integrated antigenicity score was defined by two components:

- Attention score: Only correctly predicted samples were included. In a five-fold cross-validation framework, attention weights from correctly classified pairs were aggregated and standardized using z-score normalization, resulting in a global attention score for each site.
- Information gain: For this global analysis, information gain was computed for each site. Using HI assay data, strain pairs were labeled as antigenically similar or variant. The feature for this task was defined as the pairwise amino acid state at that specific site (e.g., “G-K” compared to “G-G”). The resulting information gain score therefore quantifies the contribution of the amino acid pair at each site to distinguishing between antigenically similar and variant strain pairs.

Integrated antigenicity score were calculated for all amino acid positions in the HA1 protein, and sites were ranked by global integrated antigenicity score. The top 40 sites, with the highest scores, were defined as global key sites (Fig. 1B), reflecting their consistent influence on antigenic properties throughout influenza virus evolution.

#### Identification of cluster-transition determining sites

To identify sites driving transitions between successive antigenic clusters, we performed a targeted analysis. Antigenic cluster boundaries were first determined (see Section Antigenic cluster inference), and a subset of strain pairs spanning adjacent clusters (cluster n and cluster n+1) was constructed. Integrated antigenicity score was recalculated for this subset, with its two components defined as follows:

- Attention score: only inter-cluster pairs were considered; attention weights were summed and z-score normalized to capture sites critical for identifying cluster transitions.
- Information gain: For this cluster-transition analysis, information gain was computed for each site. The classification label was shifted from antigenic relationship to cluster membership (discriminating between cluster n and cluster n+1). The feature for this task was defined as the single amino acid state at that specific site (e.g., ‘G’ or ‘K’). The resulting information gain score therefore quantifies the discriminatory power of the amino acid state at each site for classifying a strain as belonging to either cluster n or cluster n+1.

The integrated antigenicity score ranking reflects each site’s importance in driving cluster shifts. High-ranking sites are likely cluster-transition–determining sites. To refine candidates, we intersected the top 20 globally important sites with the top 5 cluster-transition sites. Only sites meeting both criteria were considered core candidates potentially triggering antigenic cluster transitions through single mutations (Fig. 1C), providing insights for understanding and predicting viral antigenic evolution.

### Experimental design

To evaluate the performance of PREDAC-Transformer, we conducted a two-stage benchmarking experiment. First, we determined the optimal model configuration by testing three feature encoding schemes (7-features only, ESM-2 only, and the hybrid 7-features + ESM-2) and optimizing hyperparameters across ten different model architectures.

In the second stage, we compared the optimized PREDAC-Transformer against several state-of-the-art models. For influenza A/H3N2, the competitors included PREDAC-CNN (Meng et al. 2024), IAV-CNN (Yin et al. 2022), PREDAV-FluA (Peng et al. 2017), Lee (Lee and Chen 2004), Lees (Lees et al. 2010), and PREDAC-H3 (Du et al. 2012). For A/H1N1, we compared it with PREDAC-H1 (Liu et al. 2015). We employed two evaluation strategies. First, we performed 5-fold cross-validation, where the dataset was randomly split into training (80%), validation (10%), and testing (10%) sets for five independent runs. Second, we conducted retrospective testing to simulate a real-world vaccine recommendation scenario. For a given target year, models were trained on all data available up to the preceding year and tested on strain pairs from the target year. Performance was assessed using the Area Under the Receiver Operating Characteristic Curve (AUROC) and the Area Under the Precision-Recall Curve (AUPRC) as the primary evaluation metrics.

## Results

### Overview of PREDAC-Transformer

In this study, we developed PREDAC-Transformer, a model based on the Transformer encoder architecture, to predict the antigenic evolution of seasonal influenza A viruses (Figure 1). The model takes paired HA1 sequences from influenza A/H3N2 or A/H1N1 viruses as input. PREDAC-Transformer generates a hybrid input matrix by integrating the physicochemical features of amino acids with contextual embeddings from the ESM-2 language model. Using this input representation, the model predicts the antigenic relationship between paired influenza strains. Subsequently, it constructs a distance matrix from the model’s Softmax probabilities and infers antigenic clusters using UMAP for dimensionality reduction and K-means for clustering.

Leveraging the attention weights from PREDAC-Transformer, we established a multi-level framework to identify key amino acid sites driving antigenic variation. A key innovation of our work is the integrated antigenicity score, a metric designed to assess the importance of individual amino acid sites. The integrated antigenicity score integrates two components: attention scores from the Transformer’s multi-head self-attention matrix, which quantifies the degree of attention the model assigns to different sites during prediction, and information gain derived from HI assay data or clustering results, which measures a site’s contribution to distinguishing between antigenic similarity and distinct. By integrating model attribution (attention scores) with data-driven statistical significance (information gain), a site is identified as high-scoring only when it stands out in both dimensions. This effectively reduces false positives arising from model artifact, as well as frequent mutations unrelated to antigenicity. Based on the integrated antigenicity score, PREDAC-Transformer identifies two categories of key sites: global key sites, which have a sustained role in long-term antigenic evolution, and cluster-transition determining sites, which are critical for transitions between antigenic clusters.

### Performance Evaluation of PREDAC-Transformer

To systematically evaluate the performance of PREDAC-Transformer, we conducted a two-stage benchmarking experiment. The first stage assessed the impact of different feature encodings (7-features, ESM-2, and the hybrid ESM-2+7-features) and optimizing hyperparameters across ten model architectures. The second stage compared the optimized model against six state-of-the-art competitors.

In 5-fold cross-validations, as shown in the Supplementary Figures S1-S2, the hybrid ESM-2+7- features encoding yielded the best performance, highlighting the advantage of this approach. This method leverages key physicochemical properties while incorporating deep semantic information from the ESM-2 model, providing a more comprehensive input for predicting antigenic relationships. Regarding the model architecture, we performed separate optimizations for the influenza A/H3N2 and A/H1N1 viruses. We tested the impact of varying the number of encoder layers and attention heads on the performance of antigenic relationship prediction. The results demonstrated that for the A/H3N2 virus, the model achieved optimal performance with a 2-layer, 8-head architecture, yielding an AUROC of 0.992 in 5-fold cross-validation. For the A/H1N1 virus, a 2-layer, 4-head architecture proved more effective, achieving an AUROC of 0.921 in 5-fold cross-validation. This difference is likely attributable to the distinct evolutionary characteristics of their HA1 sequences. The more complex antigenic drift patterns of A/H3N2 may require more attention heads to capture subtle sequence variations (Smith et al. 2004; Bedford et al. 2014).

In comparisons with six state-of-the-art models, PREDAC-Transformer achieved the highest performance in 5-fold cross-validations (Figure 2A, 2B, Supplementary Tables S3-S6). Specifically, compared to PREDAC-CNN, the best-performing competitor, PREDAC-Transformer increased the AUC from 0.906 to 0.921 for A/H1N1 and from 0.990 to 0.992 for A/H3N2. For retrospective testing, which simulates real-world vaccine strain recommendation, models were trained on data up to the year preceding a target year and tested on that target year’s data. Across 17 annual subsets for A/H3N2 and A/H1N1, PREDAC-Transformer again outperformed the other models, with the performance gains being more pronounced (Figure 2C and 2D, Supplementary Tables S7-S8). The 2012 subset for A/H3N2 was excluded from calculations due to the absence of antigenic distinct pairs. PREDAC-Transformer improved the overall AUPRC from 0.805 to 0.822 for A/H1N1 and from 0.801 to 0.839 for A/H3N2. This substantial improvement is likely due to the Transformer architecture’s enhanced ability to capture complex dependencies and predict novel drift patterns, which is critical for the challenges of retrospective testing.

**Figure 2.**
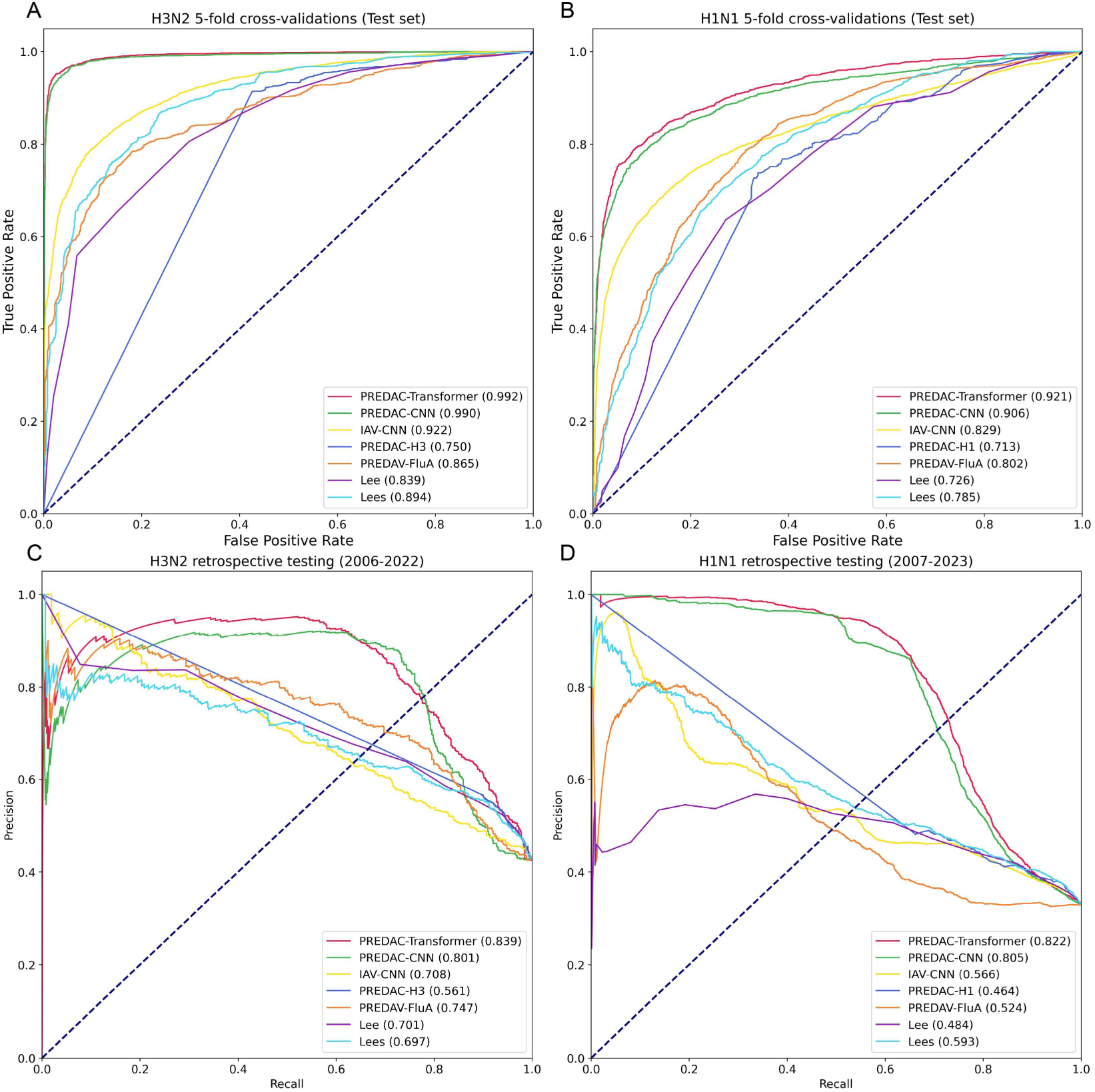
Performance comparison of PREDAC-Transformer with competitor models using 5-fold cross-validations and retrospective testing. ROC curves for 5-fold cross-validation on influenza A/H3N2 (A) and A/H1N1 (B), and Precision-Recall (PR) curves for retrospective testing on A/H3N2 (C) and A/H1N1 (D) are shown.

### An Attention-Based Perspective for Analyzing Antigenic Differences: A Single-Strain-Pair Case Study

Following the prediction of antigenic relationships, the central challenge lies in identifying the sites responsible for antigenic differences. We addressed this by analyzing the attention weight distributions of representative strain pairs to explore the decision-making process of the model. As shown in Figure 3, the model identifies not only key sites responsible for cluster transitions but also other functional regions previously reported in the literature.

**Figure 3.**
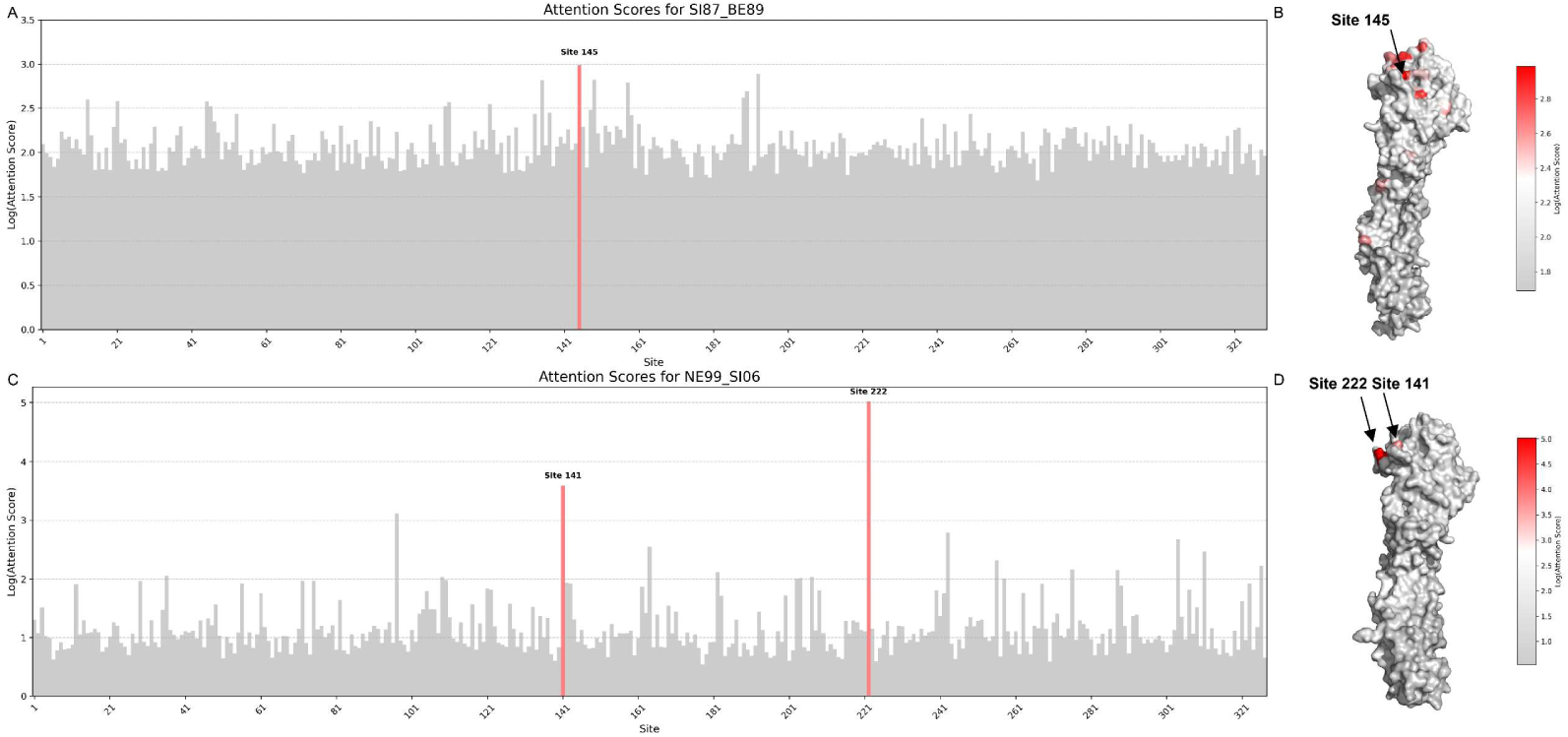
Attention weight analysis of representative strain pairs. (A, C) Bar plots displaying attention weights for each amino acid position in the HA protein of (A) Influenza A/H3N2 and (C) Influenza A/H1N1. Literature-defined antigenic sites are highlighted in red. (B, D) Attention weights mapped onto the HA protein structure for (B) H3N2 (PDB: 2YPG) and (D) H1N1 (PDB: 2WRG). Structures are colored by attention weight, progressing from gray (low) to red (high).

In the analysis of the SI87 and BE89 strain pair, a classic example of an H3N2 cluster transition, the site with the highest attention weight (position 145) corresponds to the key antigenic site identified by Koel et al (Koel et al. 2013). as driving the cluster transition. When mapped onto the HA protein’s three-dimensional structure, these high-attention sites clustered significantly within the antigenic ridge region (Figure 3A, 3B), a primary target for antibody recognition (Koel et al. 2013). Similarly, for the H1N1 subtype, the model correctly identified the cluster transition site 141 (position 140 in H1 numbering) when analyzing the NE99 and SI06 strains. It also highlighted position 222 (Figure 3C, 3D), a site known to be critical for receptor-binding specificity and interspecies transmission (Chutinimitkul et al. 2010). These results indicate that the model’s attention mechanism effectively captures functionally important sites influencing antigenicity.

Although attention weights excel at narrowing the scope of key site identification, relying solely on them presents limitations. A model can carry architectural biases: its attention heads may systematically overweight sequence edges (start/end) that lack biological relevance, conflating true signal with model artifacts (Jain and Wallace 2019; Wang et al. 2023; Mistry et al. 2025). For example, in this study, sites of the H3N2 subtype pair SI87 and BE89 near the beginning of the sequence received high attention weights which are not directly involved in receptor binding and are inconsistent with previously reported antigenic regions. This phenomenon may be attributable to a “primacy effect” within the Transformer’s attention heads (Wang et al. 2023; Mistry et al. 2025). Consequently, attention mechanisms cannot be directly equated with causal explanation (Jain and Wallace 2019). Therefore, establishing a reliable scoring system requires calibration with evidence independent of the model’s architecture to reflect true biological importance, which provided the rationale for developing the integrated antigenicity score.

### Identification and Evaluation of Global Key Sites via the Integrated Antigenicity Score

We propose a multi-level analysis framework based on the attention weights of PREDAC-Transformer to identify amino acid sites driving antigenic variation. To assess its effectiveness, we performed two evaluations. First, we validated the identified sites by comparing them with antigenicity-related sites reported in the literature, focusing on their distribution within known epitopes (five regions A–E in A/H3N2 and six regions Sa, Sb, Ca, Cb, Pa, and Pb in A/H1N1 (Quan et al. 2019) (Liu et al. 2023)). Second, we tested whether incorporating these high-scoring sites could improve the performance of an existing baseline model in antigenic relationship prediction.

To create a robust scoring system, we developed the integrated antigenicity score, which combines model attribution (attention scores) with data-driven statistical significance (information gain) to assess site importance. As shown in the Supplementary Figures S3-S4, validation experiments show that integrated antigenicity score significantly outperforms traditional metrics such as mutation rate and site entropy in identifying experimentally verified antigenicity-related sites (Mann-Whitney U, p < 0.001, Supplementary Figure S4). To evaluate the contribution of integrated antigenicity score components and validate the integrated design, we conducted ablation experiments on H1N1 and H3N2 (Supplementary Figure S3). Results indicate that integrated antigenicity score is most reliable in distinguishing antigenic sites, exhibiting the largest effect sizes. Specifically, for H1N1, Cohen’s d for attention score and information gain were 0.32 and 0.31 (small effect), respectively, while integrated antigenicity score reached 0.43 (moderate effect). For H3N2, Cohen’s d for attention score and information gain were 0.58 and 0.62 (moderate effect), respectively, while integrated antigenicity score reached 0.86 (large effect). This demonstrates that integrating attention and information gain significantly enhances the accuracy and robustness of antigenic site identification.

Based on integrated antigenicity scores, we defined the top 40 scoring sites as global key sites (Figure 4, Tables 1 and 2). The complete ranking of sites is provided in Supplementary Tables S9-S10. For the H3N2 virus, 39 of these 40 sites are located within known antigenic epitopes (A-E), and all seven “gold standard” sites for antigenic drift proposed by Koel et al. were included (Koel et al. 2013). The remaining site (225) is adjacent to epitope D and has been reported to affect receptor-binding affinity (Thompson et al. 2024). For the H1N1 virus, 19 of the top 40 sites are within its known antigenic epitopes (Sa, Sb, Ca, Cb, Pa, Pb), and 12 sites have been previously reported in the literature as being associated with antigenicity (Koel et al. 2013; Harvey et al. 2016; Lin et al. 2024; Maurer et al. 2025).The distribution of these global key sites revealed distinct patterns (Figure 4). For A/H3N2, sites were highly concentrated in epitopes B (n=12), A (n=11), and D (n=9), suggesting that its antigenic drift is driven by mutations in a few immunodominant regions. In contrast, the distribution for A/H1N1 was more dispersed, with the highest concentration in epitope Sa (n=8) and a significant number of important sites located outside of canonical epitopes.

**Figure 4.**
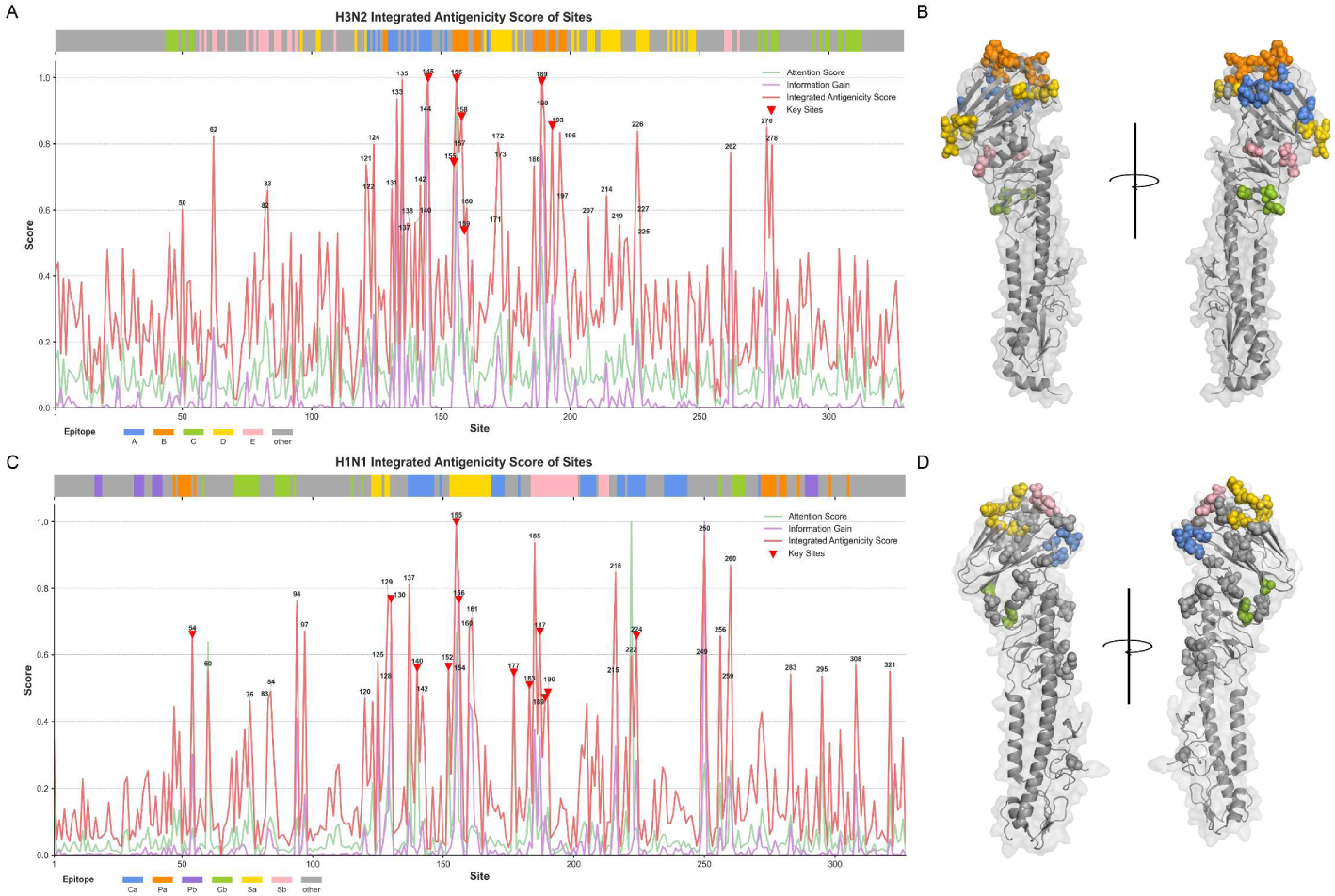
Identification and structural mapping of global key sites using the integrated antigenicity score. (A, C) Line plots comparing attention score (green), information gain (purple), and the integrated antigenicity score (red) across HA1 amino acid positions for (A) H3N2 and (C) H1N1. The top 40 sites with the highest integrated antigenicity score, designated as global key sites, are labeled with their corresponding site numbers. Known epitope regions are annotated above the plots, and literature-defined antigenic sites are marked with red inverted triangles for reference. (B, D) Structural mapping of global key sites onto the HA protein for (B) H3N2 (PDB: 2YPG) and (D) H1N1 (PDB: 2WRG). Identified sites are color-coded by their corresponding epitope region.

**Table 1.** Integrated antigenicity score assessment of top 40 in H3N2. information_gain.

| Rank | Position | Score_attention | Score_information_gain | Integrated antigenicity score | Epitope |
| --- | --- | --- | --- | --- | --- |
| 1 | 145 | 97222.945 | 0.301 | -0.001 | A |
| 2 | 156 | 89510.590 | 0.215 | -0.005 | B |
| 3 | 135 | 65573.875 | 0.253 | -0.016 | A |
| 4 | 189 | 63632.676 | 0.239 | -0.022 | B |
| 5 | 133 | 50700.918 | 0.206 | -0.144 | A |
| 6 | 144 | 49552.676 | 0.099 | -0.257 | A |
| 7 | 158 | 49313.160 | 0.099 | -0.263 | B |
| 8 | 190 | 45890.816 | 0.184 | -0.284 | B |
| 9 | 193 | 46816.938 | 0.103 | -0.326 | B |
| 10 | 276 | 45525.418 | 0.123 | -0.339 | C |
| 11 | 226 | 49260.297 | 0.071 | -0.363 | D |
| 12 | 196 | 49589.750 | 0.067 | -0.373 | B |
| 13 | 62 | 47604.660 | 0.073 | -0.397 | E |
| 14 | 172 | 47328.098 | 0.065 | -0.442 | D |
| 15 | 124 | 44834.836 | 0.085 | -0.452 | A |
| 16 | 278 | 46663.824 | 0.067 | -0.457 | C |
| 17 | 157 | 41881.860 | 0.127 | -0.509 | B |
| 18 | 262 | 41134.598 | 0.218 | -0.512 | E |
| 19 | 173 | 50328.023 | 0.037 | -0.566 | D |
| 20 | 155 | 46773.710 | 0.047 | -0.573 | B |
| 21 | 121 | 44835.090 | 0.055 | -0.593 | D |
| 22 | 186 | 45538.914 | 0.050 | -0.599 | B |
| 23 | 142 | 42787.477 | 0.049 | -0.734 | A |
| 24 | 131 | 44308.285 | 0.037 | -0.758 | A |
| 25 | 83 | 46755.540 | 0.026 | -0.764 | E |
| 26 | 122 | 47364.934 | 0.023 | -0.786 | A |
| 27 | 214 | 42773.540 | 0.040 | -0.803 | D |
| 28 | 197 | 40225.695 | 0.055 | -0.847 | B |
| 29 | 160 | 47417.387 | 0.014 | -0.885 | B |
| 30 | 50 | 41713.926 | 0.036 | -0.895 | C |
| 31 | 82 | 49664.210 | 0.008 | -0.913 | E |
| 32 | 207 | 49387.594 | 0.006 | -0.946 | D |
| 33 | 227 | 42564.637 | 0.024 | -0.978 | D |
| 34 | 140 | 45875.188 | 0.011 | -0.987 | A |
| 35 | 137 | 38884.945 | 0.048 | -0.991 | A |
| 36 | 138 | 48571.367 | 0.004 | -0.996 | A |
| 37 | 219 | 43918.800 | 0.016 | -1.002 | D |
| 38 | 171 | 46044.297 | 0.009 | -1.009 | D |
| 39 | 225 | 40700.000 | 0.030 | -1.018 | Other (Literature Reported) |
| 40 | 159 | 38226.620 | 0.048 | -1.039 | B |

**Table 2.** Integrated antigenicity score assessment of top 40 in H1N1. information_gain.

| Rank | Position | Score_attention | Score_information_gain | Integrated antigenicity score | Epitope |
| --- | --- | --- | --- | --- | --- |
| 1 | 155 | 79845.210 | 0.073 | -0.027 | Sa |
| 2 | 250 | 42158.105 | 0.172 | -0.101 | Other |
| 3 | 185 | 41610.020 | 0.064 | -0.146 | Sb |
| 4 | 260 | 42901.330 | 0.035 | -0.274 | Other |
| 5 | 216 | 32982.324 | 0.056 | -0.313 | Other |
| 6 | 137 | 53610.832 | 0.022 | -0.382 | Ca |
| 7 | 129 | 41506.600 | 0.023 | -0.439 | Sa |
| 8 | 130 | 26411.084 | 0.110 | -0.462 | Other (Literature Reported) |
| 9 | 156 | 26238.418 | 0.147 | -0.466 | Sa |
| 10 | 94 | 26799.137 | 0.071 | -0.472 | Other |
| 11 | 161 | 24295.188 | 0.075 | -0.572 | Sa |
| 12 | 160 | 22896.186 | 0.078 | -0.634 | Sa |
| 13 | 97 | 27371.758 | 0.031 | -0.646 | Other |
| 14 | 187 | 23207.809 | 0.061 | -0.648 | Sb |
| 15 | 54 | 23475.310 | 0.052 | -0.664 | Other (Literature Reported) |
| 16 | 224 | 23607.453 | 0.049 | -0.672 | Ca |
| 17 | 256 | 36636.504 | 0.014 | -0.674 | Cb |
| 18 | 222 | 112203.945 | 0.003 | -0.787 | Ca |
| 19 | 249 | 29992.514 | 0.015 | -0.801 | Other |
| 20 | 125 | 19626.357 | 0.075 | -0.816 | Sa |
| 21 | 308 | 21124.512 | 0.042 | -0.839 | Other |
| 22 | 152 | 49900.617 | 0.002 | -0.845 | Other (Literature Reported) |
| 23 | 140 | 67351.340 | 0.001 | -0.852 | Ca |
| 24 | 60 | 77353.000 | 0.000 | -0.867 | Other |
| 25 | 321 | 33289.945 | 0.008 | -0.873 | Other |
| 26 | 177 | 63548.992 | 0.000 | -0.879 | Other (Literature Reported) |
| 27 | 154 | 28810.270 | 0.013 | -0.887 | Sa |
| 28 | 283 | 27374.883 | 0.015 | -0.889 | Other |
| 29 | 295 | 45512.336 | 0.001 | -0.899 | Other |
| 30 | 215 | 28316.697 | 0.013 | -0.903 | Other |
| 31 | 128 | 23782.633 | 0.021 | -0.934 | Sa |
| 32 | 259 | 19592.480 | 0.040 | -0.936 | Other |
| 33 | 183 | 32041.098 | 0.007 | -0.949 | Other (Literature Reported) |
| 34 | 84 | 26789.580 | 0.012 | -0.983 | Other |
| 35 | 190 | 29719.738 | 0.008 | -0.990 | Sb |
| 36 | 142 | 23552.941 | 0.018 | -1.008 | Ca |
| 37 | 189 | 22267.643 | 0.020 | -1.022 | Sb |
| 38 | 120 | 24842.922 | 0.014 | -1.024 | Other |
| 39 | 83 | 24810.875 | 0.014 | -1.030 | Other |
| 40 | 76 | 36971.780 | 0.000 | -1.039 | Cb |

Structurally, for both subtypes, most important sites are located in the receptor-binding domain (RBD) at the head of the HA protein. In A/H3N2, these sites correspond to highly exposed loops on the HA surface that form the antigenic ridge. In A/H1N1, key sites were found not only in the RBD but also in the vestigial esterase domain (ED) (Gamblin et al. 2021), suggesting a potentially unique immune evasion mechanism involving conformational changes induced by mutations outside the RBD. For example, some broad-spectrum antibodies targeting the vestigial esterase subdomain can affect the binding of the virus to the receptor by inhibiting membrane fusion (Iba et al. 2014; Jiao et al. 2023).

To assess the practical value of the identified high-scoring sites in antigenic relationship prediction tasks, we designed an experiment using PREDAC-CNN as a benchmark model. To test whether focusing on key antigenic determinants improves predictive performance, we modified the PREDAC-CNN model by increasing its attention on specific site sets. We compared the performance of this weighted model when using the four different input feature sets: the full-length HA1 sequence, traditional antigenic epitopes, the top 20 integrated antigenicity score-ranked sites, and the top 40 integrated antigenicity score-ranked sites, which correspond to our definition of global key sites (Supplementary Figure S5). Results showed that the top 40 high-scoring sites achieved the best performance across all test subsets, validating their suitability as a core feature set. The performance improvement was particularly pronounced in the more challenging retrospective tests. The AUPRC for A/H3N2 viruses increased significantly from 0.801 to 0.857, while for A/H1N1 viruses, it improved from 0.805 to 0.814. In 5-fold cross-validation, although the baseline model’s performance on the full-length sequence (H3: 0.990, H1: 0.906) was already approaching saturation, using the top 40 sites delivered stable and optimal performance (H3: 0.991, H1: 0.908), outperforming the top 20 sites as well. This demonstrates that the integrated antigenicity score can accurately identify the core sites critical for antigenic relationship prediction from the full sequence, and that the top 40 size achieves an optimal balance between retaining key information and filtering out noise.

### Identification of Historical Cluster-Transition Determining Sites

The antigenic evolution of influenza viruses occurs in a cluster-wise manner, leading to distinct antigenic landscape in different seasons. Identifying these antigenic clusters and the key sites driving transitions between them is essential for understanding viral evolutionary pathways and guiding vaccine strain selection. We systematically classified antigenic clusters for A/H3N2 viruses from 1968 to 2024 and A/H1N1 viruses from 1977 to 2024, identifying 24 and 12 major clusters, respectively (Tables 3-4, Supplementary Figures S7-S8). The identified clusters and their circulation patterns are highly consistent with those reported in previous studies, supporting the reliability of our methodology (Smith et al. 2004; Liu et al. 2023; Cheng et al. 2024). The majority of vaccine strains within each antigenic cluster were consistent with HI titration data. However, we also observed that a few strains, although assigned to a primary cluster, exhibited partial antigenic similarity with strains from other clusters, reflecting the intrinsic networked and continuous characteristics of the antigenic space. For instance, the A/H3N2 strain A/Darwin/9/2021 (DA21 cluster) is antigenically similar to A/Thailand/8/2022 and A/Massachusetts/18/2022 (MA22 cluster). Similarly, in H1N1, A/Victoria/2570/2019 (VI19 cluster) and A/Sydney/5/2021 (SY21 cluster) share antigenic similarity. Our algorithm assigned such strains to the cluster of highest overall similarity, which aligns with WHO vaccine recommendation practices. Building on this consistency, we analyzed antigenic cluster dynamics up to May 2024 to further demonstrate the framework’s utility for vaccine strain recommendation. We assessed whether dominant clusters matched vaccine strains and tracked emerging variants (Supplementary Figure S9). As of February 2024, the dominant cluster remained aligned with the vaccine strain, consistent with WHO’s 2025 recommendations. For the September 2025 Southern Hemisphere recommendations (unpublished, Supplementary Figure S10), we identified a new minor H3N2 cluster (subclade J.2.4, T135K in epitope A, ranked third in global key sites) that expanded in late 2024, and the strain within this cluster (A/Singapore/GP20238/2024) was subsequently confirmed to be consistent with the recent Southern Hemisphere recommended vaccine strains updated by the WHO for use in 2026 (Anon). Accurate delineation of antigenic clusters facilitates vaccine strain recommendation, and pinpointing cluster-transition sites is equally crucial for narrowing surveillance targets and refining candidate vaccine selection.

**Table 3.** Summary of 25 predicted antigenic clusters for influenza A (H3N2) viruses.

| Cluster | Circulation time | Strain number | World Health Organization (WHO) recommended vaccine strains |
| --- | --- | --- | --- |
| HK68 | 1968-1973 | 64 | A/Hong_Kong/1/1968 |
| EN72 | 1972-1977 | 47 | A/England/42/1972 |
| VI75 | 1975-1978 | 30 | A/Victoria/3/1975 |
| TX77 | 1976-1987 | 93 | A/Texas/1/1977, A/Bangkok/1/1979, A/Philippines/2/1982, A/Christchurch/4/1985, A/Mississippi/1/1985 |
| BK79 | 1978-1981, 1992-1996, 2005-2024 | 3640 | A/Bangkok/1/1979, A/Beijing/32/1992, A/Moscow/10/1999 |
| SI87 | 1987-1992 | 55 | A/Shanghai/11/1987, A/Sichuan/2/1987, A/Guizhou/54/1989 |
| BE89 | 1988-1994 | 102 | A/Beijing/353/1989 |
| BE92 | 1991-1997 | 77 | A/Beijing/32/1992, A/Shangdong/9/1993, A/Johannesburg/33/1994 |
| WU95 | 1993-1999 | 93 | A/Wuhan/359/1995 |
| SY97 | 1996-2006, 2013-2016 | 420 | A/Sydney/5/1997, A/Moscow/10/1999 |
| FU02 | 2002-2008 | 731 | A/Fujian/411/2002, A/Wellington/1/2004 |
| CA04 | 2003-2013 | 2162 | A/California/7/2004, A/Wisconsin/67/2005, A/Brisbane/10/2007 |
| PE09 | 2009-2014 | 619 | A/Perth/16/2009 |
| VI11 | 2009-2018 | 6866 | A/Victoria/361/2011, A/Texas/50/2012 |
| SW13 | 2013-2020 | 1113 | A/Switzerland/9715293/2013 |
| HK14 | 2013-2019 | 9683 | A/Hong_Kong/4801/2014 |
| SG16 | 2014-2021 | 8354 | A/Singapore/INFIMH-16-0019/2016 |
| KS17 | 2014-2022 | 7338 | A/Kansas/14/2017 |
| SW17 | 2014-2020 | 10104 | A/Switzerland/8060/2017 |
| HK19 | 2017-2023 | 1807 | A/Hong_Kong/2671/2019, A/Hong_Kong/45/2019 |
| SA19 | 2016-2022 | 9246 | A/South_Australia/34/2019 |
| CA20 | 2018-2024 | 12021 | A/Cambodia/e0826360/2020 |
| DA21 | 2020-2024 | 18702 | A/Darwin/6/2021, A/Darwin/9/2021 |
| MA22 | 2021-2025 | 6512 | A/Thailand/8/2022, A/Massachusetts/18/2022 |
| DC23 | 2020-2025 | 12914 | A/District_Of_Columbia/27/2023, A/Croatia/10136RV/2023 |

**Table 4.** Summary of 12 predicted antigenic clusters for influenza A (H1N1) viruses.

| Cluster | Circulation time | Strain number | World Health Organization (WHO) recommended vaccine strains |
| --- | --- | --- | --- |
| US77 | 1977-1984,<br>1990-1993,<br>2020-2023 | 116 | A/USSR/90/1977, A/Brazil/11/1978,<br>A/Chile/1/1983 |
| SI86 | 1986-1990,<br>1991-1994,<br>1995-1999 | 90 | A/Singapore/6/1986, A/Bayern/7/1995 |
| BE95 | 1995-2009 | 860 | A/Beijing/262/1995 |
| NE99 | 1999-2009 | 120 | A/New_Caledonia/20/1999 |
| SI06 | 2005-2010 | 220 | A/Solomon_Islands/03/2006 |
| BR07 | 2006-2011 | 764 | A/Brisbane/59/2007 |
| CA09 | 2009-2017 | 12765 | A/California/7/2009 |
| MI15 | 2007-2025 | 15505 | A/Michigan/45/2015 |
| BR18 | 2008-2022 | 13411 | A/Brisbane/02/2018 |
| VI19 | 2008-2025 | 3605 | A/Victoria/2570/2019, A/Wisconsin/588/2019 |
| GD19 | 2018-2025 | 11204 | A/Guangdong-Maonan/SWL1536/2019,<br>A/Hawaii/70/2019 |
| SY21 | 2021-2025 | 30862 | A/Sydney/5/2021, A/Victoria/4897/2022,<br>A/Wisconsin/67/2022 |

We have mapped the major antigenic clusters, revealing that influenza’s antigenic evolution follows a pattern characterized by distinct clusters. While global key sites play a role in overall antigenicity, a critical subsequent question is to find which sites are specifically responsible for driving the transitions between these clusters. To identify the most critical sites driving these transitions while minimizing noise, we implemented a rigorous selection strategy for cluster-transition determining sites. Based on the characteristics of established “gold standard” sites, we required a site to meet two criteria: global importance across evolutionary history (ranking in the top 20 for global integrated antigenicity score) and specificity to cluster transitions (ranking in the top 5 for inter-cluster integrated antigenicity score). Sites that satisfied both criteria were designated as cluster-transition determining sites.

Our results show that 17 cluster-transition determining sites were involved in 27 cluster transitions for A/H3N2, while 14 cluster-transition determining sites were involved in 12 transitions for A/H1N1 (Figure 5 and Supplementary Tables S11-S12). The complete ranking of sites is provided in Supplementary Tables S13-S14. The high enrichment of cluster-transition determining sites within functionally important and antigenically relevant regions provides biological validation for our approach. The majority of cluster-transition determining sites reside within canonical antigenic epitopes, underscoring their critical role in antigenic drift. Furthermore, several cluster-transition determining sites are located at known glycosylation sites (e.g., positions 135, 144, and 145 in H3N2; 54, 125, 129, and 130 in H1N1 (Tate et al. 2014; Huang et al. 2017; Li et al. 2020)), suggesting they may influence antigenicity by altering surface glycan shields. Notably, our method identified functionally significant sites previously reported in the literature, such as positions 133, 135, 145, 155, 156, 158, 172, 173, and 193 in H3N2, which are known broad-spectrum antibody epitopes (Lee et al. 2014). All the seven cluster transition sites experimentally validated by Koel et al. (Koel et al. 2013) were included among our identified cluster-transition determining sites, further confirming the accuracy of our method. For A/H1N1, a single mutation at position 156 is known to cause a 4-fold to 8-fold reduction in HI titers (Li et al. 2016), and sites 125, 129, 187, and 224 have also been reported to affect antigenicity (Koel et al. 2015; Li et al. 2016; Lin et al. 2024; Swanson et al. 2024). These findings affirm the critical role of the identified sites in viral immune escape.

**Figure 5.**
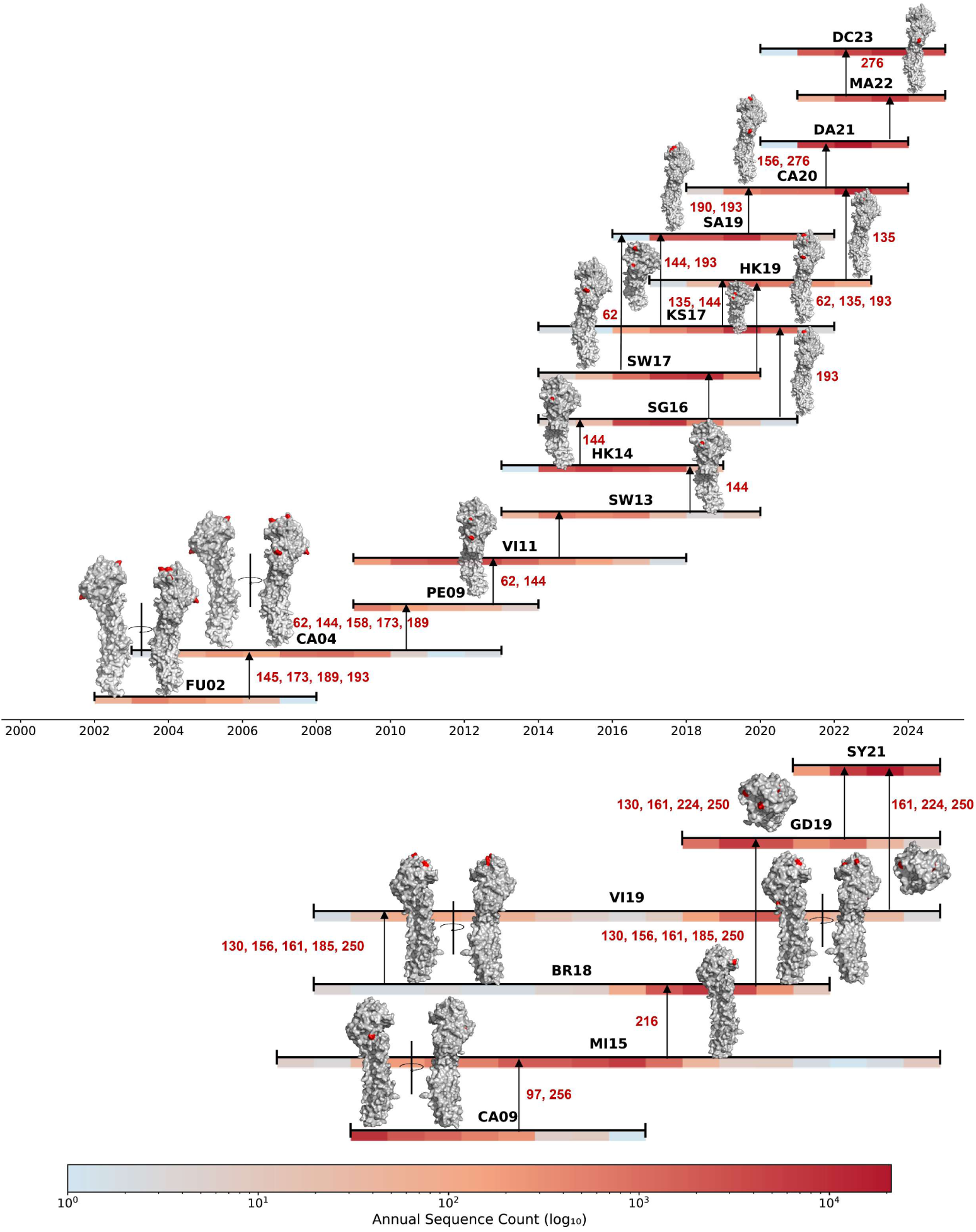
Cluster-transition determining sites in the antigenic evolution of influenza. A viruses. The figure illustrates antigenic cluster evolution since 2000 (the full version is provided in Supplementary Tables S11 and S12), with A/H3N2 shown on the top and A/H1N1 on the bottom. Each cluster is represented by a hemagglutinin (HA) trimer structure of H3N2 (PDB: 2YPG) and H1N1 (PDB: 2WRG), where cluster-transition determining sites are highlighted in red and annotated with their sequence positions. Horizontal heatmaps beneath each structure indicate the circulation period of each cluster, with color intensity (blue to red, log scale) reflecting annual isolate counts. Arrows denote the major evolutionary directions.

In summary, the dynamic monitoring of antigenic clusters with PREDAC-Transformer provides timely and quantitative data to support global influenza surveillance and vaccine decision-making. Moreover, the identification of cluster-transition determining sites enables rapid, low-cost monitoring of potential antigenic changes at the molecular level. Integrating this information into surveillance systems can help narrow the scope of vaccine candidate screening and enhance the precision of influenza prevention and control.

## Discussion

Accurately predicting the antigenic relationships of influenza viruses and identifying the key features that drive antigenic drift are pivotal for vaccine strain recommendation. Previous approaches relying on known epitopes limit exploration of complex site interactions. Deep learning methods like CNNs lack interpretability, making it hard to understand their predictions, and struggle to capture long-range dependencies essential for antigenicity. To address these limitations, we developed PREDAC-Transformer. This model not only surpasses existing benchmark methods in predicting antigenic variants but also demonstrates superior performance in both 5-fold cross-validation and retrospective testing, validating its effectiveness for influenza antigenic relationship prediction. Using the model, antigenic clusters can be inferred more accurately, enabling high-resolution analysis of the extent of antigenic cluster changes. More importantly, PREDAC-Transformer introduces a novel analytical framework that spans from prediction to interpretation, offering a new perspective for understanding the mechanisms of influenza’s antigenic evolution. In this study, we convert the model’s internal attention weights into a quantifiable metric, the integrated antigenicity score. The integrated antigenicity score reflects the sequence signals that the model prioritizes during prediction and, through an information-theoretic framework, ensures these signals are statistically correlated with antigenic differences. Based on the integrated antigenicity score, we successfully identified global key sites and cluster-transition determining sites, providing new insights and methodologies to optimize vaccine strain recommendation strategies.

The strong performance of PREDAC-Transformer arises from its architecture’s consistency with the mechanisms of antibody recognition. Antigen-antibody binding is determined by the three-dimensional conformation of the antigen surface (Murin et al. 2019). For instance, the receptor-binding site (RBS) on the hemagglutinin (HA) head is formed by the spatial arrangement of multiple discontinuous secondary structures, while the canonical antigenic sites are composed of distinct loops and helices (Gamblin et al. 2021). This structural complexity presents a fundamental challenge for prediction models based solely on the primary sequence. Such complexity challenges sequence-based methods. Traditional CNNs have been widely used but come with certain limitations (Meng et al. 2021; Meng et al. 2024). For instance, they lack interpretability, making it difficult to understand the underlying mechanisms of their predictions. Furthermore, CNNs are primarily designed to capture local features and struggle with identifying long-range dependencies that are crucial for conformational epitopes (Murin et al. 2019; Yuan et al. 2023). In contrast, the self-attention mechanism of the Transformer architecture models global dependencies by calculating association weights between all amino acid pairs in a sequence. This not only enhances its ability to capture long-range interactions but also provides interpretability, enabling researchers to identify the key sequence regions driving predictions. These strengths make the Transformer a more powerful tool for antigenic relationship prediction based on primary sequences. Building on this architecture, we introduced a multimodal feature fusion strategy that integrates raw sequences, physicochemical properties, and ESM-2 embeddings derived from a large-scale protein language model. By integrating sequence, biophysics, and evolutionary context, the model builds biologically relevant representations that enhance performance, especially in retrospective tests requiring generalization to novel drift patterns. Compared to the state-of-the-art PREDAC-CNN, PREDAC-Transformer increased the AUPRC from 0.805 to 0.822 for A/H1N1 viruses and from 0.801 to 0.839 for A/H3N2 viruses. These gains highlight PREDAC-Transformer’s superior ability to model complex interactions and deliver more accurate, robust predictions.

Then, we systematically inferred antigenic clusters for A/H3N2 (1968–2024) and A/H1N1 (1977–2024) viruses, identifying 24 and 12 major clusters, respectively. The temporal distribution and transition points of these clusters align well with previous studies, confirming that our model accurately captures historical evolutionary dynamics (Smith et al. 2004; Liu et al. 2023; Cheng et al. 2024). These findings provide a robust foundation for influenza surveillance and vaccine strain recommendation. The global key sites identified by the integrated antigenicity score can be used as prior information to guide and optimize other computational models, helping them focus on the most critical residues to improve prediction accuracy. Furthermore, our framework provides critical decision support for vaccine strain selection. By dynamically monitoring the antigenic match between circulating strains and current vaccine strains, we can provide timely computational evidence to inform whether a vaccine update is necessary. Our analysis of global influenza dynamics indicated that the dominant circulating cluster in February 2024 remained antigenically similar to the vaccine strain, a conclusion consistent with the subsequent WHO recommendation for the 2025 Northern Hemisphere vaccine(Anon). In our analysis to support the September 2025 Southern Hemisphere recommendation, we observed that while the dominant H3N2 cluster was still DC23, an emerging sublineage (J.2.4) showed a rapid increase in prevalence since late 2024, characterized by a T135K mutation in epitope A. Site 135 ranked third in our global integrated antigenicity score importance list, suggesting the T135K mutation is highly likely to cause significant antigenic drift. This finding highlights the need for continued surveillance of this sublineage and demonstrates how our framework can narrow the selection of vaccine candidates by identifying high-risk strains and mutations.

PREDAC-Transformer learns complex patterns related to antigenicity directly from raw sequence data, enabling it to move beyond validating existing knowledge and toward discovering new biological insights. The integrated antigenicity score, derived from the model’s attention mechanism, represents a new, data-driven approach for identifying key antigenic sites. Based on the attention mechanism, the integrated antigenicity score proposed in this study provides a novel, data-driven method for identifying key sites. Compared with previous approaches, such as those based on information gain and Shannon entropy (Cheng et al. 2024; Zhai et al. 2024) or those based on the contribution of traditional machine learning models (Han et al. 2019; Quan et al. 2019; Shah et al. 2024), integrated antigenicity score innovatively combines the model’s attention mechanism with information gain. This ensures that the identified signals are not only prioritized by the model for prediction but are also statistically correlated with antigenic variation, thereby filtering out noise from model artifact or random mutations. Using integrated antigenicity score, we systematically identified key amino acid sites influencing antigenicity in H3N2 and H1N1 subtypes. Our analysis confirmed previously established sites. For instance, 39 of the 40 globally important sites we identified in H3N2 are located within known antigenic epitopes, with a high concentration in epitopes A and B, consistent with previous research (Koel et al. 2013; Shah et al. 2024). We also identified novel sites beyond canonical epitopes, suggesting that influenza antigenicity may be modulated not only by direct epitope changes but also by mechanisms such as glycosylation or synergistic interactions (Murin et al. 2019). For example, site 225 in H3N2, while not in a defined epitope, is adjacent to epitope D and has been reported to affect receptor-binding affinity (Thompson et al. 2024). In H1N1, five such sites were found, including documented glycosylation sites (54, 130, 177) known to mask epitopes and mediate immune escape (Tate et al. 2014; Li et al. 2020), as well as nearby residues (152, 183) that may affect antigenicity through epitope interactions (Koel et al. 2015; Li et al. 2016). These findings indicate that antigenic epitopes may extend beyond current definitions, with the identified sites serving as valuable additions to existing annotations. We also applied the integrated antigenicity score framework to identify cluster-transition sites. To capture the evolutionary basis of antigenic cluster transition, we focused our calculations on strain pairs from adjacent clusters, avoiding the signal dilution inherent in global averaging. We identified the 7 key amino acid substitution positions for A/H3N2 antigenic changes proposed by Koel et al. (Koel et al. 2013), confirming the method’s reliability. We further expanded this analysis to map cluster-transition sites after 2000 and visualized their quantitative weights in a structural context. In summary, the integrated antigenicity score framework not only corroborates known antigenic determinants but also uncovers novel candidate sites, offering new insights into the mechanisms of antigenic drift.

Although this study represents significant progress, several limitations remain. The model’s performance is constrained by the quantity and quality of available antigenic data. Furthermore, our model currently considers only the HA sequence, while the NA protein is also known to influence antigenicity (Wu and Ellebedy 2024). Future work will aim to incorporate additional genetic information to create a more comprehensive model. While our computational framework shows great potential, further experimental validation is needed to confirm its utility in public health decision-making. We will focus on refining the model through enhanced integration of computational and experimental approaches to improve its accuracy and reliability in practical applications.

## Authors’ contributions

Conceptualization: TJ, JM, JL

Methodology: JL, JW, CW, JM

Formal Analysis: JL, JM

Investigation: JL, JM, JW, CW, DX, WX, LQ, AW, TJ

Data Curation: JL

Writing – original draft: JL, JM

Writing – review & editing: TJ, JM, AW, XD, JW

Visualization: JL, JW

Supervision and Project administration: TJ, JM, AW

Funding acquisition and Resources: TJ, AW, XD

## Acknowledgements

This project was financially supported by the Guangzhou Laboratory Major Talent Project grant MP-GZNL2025C03010-01 (T.J.), the CAMS Innovation Fund for Medical Sciences (CIFMS) (2021-I2M-1-061, 2023-PT330-01), the National Natural Science Foundation of China (32370703) and the Suzhou Applied Basic Research Program (General Program in Medical and Health Sciences) (SYW2024065); We would like to acknowledge the support of High-throughput Sequencing and High-performance Computing Platform of Institute of Systems Medicine, Chinese Academy of Medical Sciences /Suzhou Institute of Systems Medicine (ISM).

## Data availability

Source code of PREDAC-Transformer can be accessed on Zenodo at http://doi.org/10.5281/zenodo.17421969.

