## SupplementaryMaterial for "Deciphering the Antigenic Evolution of Seasonal Influenza A Viruses with PREDAC-Transformer: From Antigenic Clustering to Key Site Identification"

* Corresponding author

**Data Collection**

**HA1 sequence of influenza A/H3N2 and A/H1N1 viruses**

(1) NCBI Influenza Virus Database (http://www.ncbi.nlm.nih.gov/genomes/FLU/Database/)

(2) Global Initiative on Sharing All Influenza Data (<http://platform.gisaid.org/>)

**HI assay data of influenza A/H3N2 and A/H1N1 viruses**

(1) Organization: Public Health Surveillance from for New Zealand.

Access: https://surv.esr.cri.nz/PDF_surveillance/Virology/FluVac/

Detail:

Recommendation for the influenza vaccine composition 2014

Recommendation for the influenza vaccine composition 2015

Recommendation for the influenza vaccine composition 2016

Recommendation for the influenza vaccine composition 2018

Recommendation for the influenza vaccine composition 2019

Recommendation for the influenza vaccine composition 2020

Recommendation for the influenza vaccine composition 2021

Recommendation for the influenza vaccine composition 2022

Recommendation for the influenza vaccine composition 2023

(2) Organization: European Centre for Disease Prevention and Control.
    Access: http://www.ecdc.europa.eu/en/PUBLICATIONS/surveillance_reports/Pages/index.aspx

Detail:

Influenza virus characterization. Summary Europe, February 2014

Influenza virus characterization. Summary Europe, March 2014

Influenza virus characterization. Summary Europe, April 2014

Influenza virus characterization. Summary Europe, May 2014

Influenza virus characterization. Summary Europe, June 2014

Influenza virus characterization. Summary Europe, July 2014

Influenza virus characterization. Summary Europe, September 2014

Influenza virus characterization. Summary Europe, November 2014

Influenza virus characterization. Summary Europe, December 2014

Influenza virus characterization. Summary Europe, February 2015

Influenza virus characterization. Summary Europe, March 2015

Influenza virus characterization. Summary Europe, April 2015

Influenza virus characterization. Summary Europe, May 2015

Influenza virus characterization. Summary Europe, June 2015

Influenza virus characterization. Summary Europe, July 2015

Influenza virus characterization. Summary Europe, September 2015

Influenza virus characterization. Summary Europe, November 2015

Influenza virus characterization. Summary Europe, December 2015

Influenza virus characterization. Summary Europe, February 2016

Influenza virus characterization. Summary Europe, March 2016

Influenza virus characterization. Summary Europe, May 2016

Influenza virus characterization. Summary Europe, June 2016

Influenza virus characterization. Summary Europe, July 2016

Influenza virus characterization. Summary Europe, September 2016

Influenza virus characterization. Summary Europe, July 2017

Influenza virus characterization. Summary Europe, September 2017

Influenza virus characterization. Summary Europe, November 2017

Influenza virus characterization. Summary Europe, December 2017

Influenza virus characterization. Summary Europe, March 2018

Influenza virus characterization. Summary Europe, May 2018

Influenza virus characterization. Summary Europe, June 2018

Influenza virus characterization. Summary Europe, September 2018

Influenza virus characterization. Summary Europe, October 2018

Influenza virus characterization. Summary Europe, November 2018

Influenza virus characterization. Summary Europe, December 2018

Influenza virus characterization. Summary Europe, March 2019

Influenza virus characterization. Summary Europe, April 2019

Influenza virus characterization. Summary Europe, May 2019

Influenza virus characterization. Summary Europe, June 2019

Influenza virus characterization. Summary Europe, September 2019

Influenza virus characterization. Summary Europe, October 2019

Influenza virus characterization. Summary Europe, November 2019

Influenza virus characterization. Summary Europe, December 2019

Influenza virus characterization. Summary Europe, February 2020

Influenza virus characterization. Summary Europe, March 2020

Influenza virus characterization. Summary Europe, April 2020

Influenza virus characterization. Summary Europe, May 2020

Influenza virus characterization. Summary Europe, June 2020

Influenza virus characterization. Summary Europe, July 2020

Influenza virus characterization. Summary Europe, September 2020

Influenza virus characterization. Summary Europe, October 2020

Influenza virus characterization. Summary Europe, November 2020

Influenza virus characterization. Summary Europe, December 2020

Influenza virus characterization. Summary Europe, February 2021

Influenza virus characterization. Summary Europe, March 2021

Influenza virus characterization. Summary Europe, May 2021

Influenza virus characterization. Summary Europe, April 2021

Influenza virus characterization. Summary Europe, June 2021

(3) Organization: U.S. Food and Drug Administration.

Access:

http://www.fda.gov/AdvisoryCommittees/CommitteesMeetingMaterials/BloodVaccinesandOtherBiologics/VaccinesandRelatedBiologicalProductsAdvisoryCommittee/default.htm.

Detail:

Information for the Vaccines and Related Biological Products Advisory Committee. March 9, 2017
Information for the Vaccines and Related Biological Products Advisory Committee. October 4, 2017
Information for the Vaccines and Related Biological Products Advisory Committee. March 1, 2018
Information for the Vaccines and Related Biological Products Advisory Committee. October 3, 2018
Information for the Vaccines and Related Biological Products Advisory Committee. March 5, 2021

Information for the Vaccines and Related Biological Products Advisory Committee. September 30, 2021

Information for the Vaccines and Related Biological Products Advisory Committee. March 3, 2022

Information for the Vaccines and Related Biological Products Advisory Committee. October 6, 2022

Information for the Vaccines and Related Biological Products Advisory Committee. March 7, 2023

Information for the Vaccines and Related Biological Products Advisory Committee. October 5, 2023

(4) Organization: World Health Organization.

Access: <http://www.who.int/wer/en/>

Detail:

Weekly Epidemiological Record 2016 No.1 ~ No.51-52

Weekly Epidemiological Record 2017 No.1 ~ No.51-52

Weekly Epidemiological Record 2018 No.1 ~ No.51-52

Weekly Epidemiological Record 2019 No.1 ~ No.51-52

Weekly Epidemiological Record 2020 No.1 ~ No.51-52

Weekly Epidemiological Record 2021 No.1 ~ No.51-52

Recommended composition of influenza virus vaccines for use in the 2015-2016 northern hemisphere influenza season

Recommended composition of influenza virus vaccines for use in the 2016 southern hemisphere influenza season

Recommended composition of influenza virus vaccines for use in the 2016-2017 northern hemisphere influenza season

Recommended composition of influenza virus vaccines for use in the 2017 southern hemisphere influenza season

Recommended composition of influenza virus vaccines for use in the 2017-2018 northern hemisphere influenza season

Recommended composition of influenza virus vaccines for use in the 2018 southern hemisphere influenza season

Recommended composition of influenza virus vaccines for use in the 2018-2019 northern hemisphere influenza season

Recommended composition of influenza virus vaccines for use in the 2019 southern hemisphere influenza season

Recommended composition of influenza virus vaccines for use in the 2019-2020 northern hemisphere influenza season

Recommended composition of influenza virus vaccines for use in the 2020 southern hemisphere influenza season

Recommended composition of influenza virus vaccines for use in the 2020-2021 northern hemisphere influenza season

Recommended composition of influenza virus vaccines for use in the 2021 southern hemisphere influenza season

Recommended composition of influenza virus vaccines for use in the 2021-2022 northern hemisphere influenza season

Recommended composition of influenza virus vaccines for use in the 2022 southern hemisphere influenza season

Recommended composition of influenza virus vaccines for use in the 2022-2023 northern hemisphere influenza season

Recommended composition of influenza virus vaccines for use in the 2023 southern hemisphere influenza season

Recommended composition of influenza virus vaccines for use in the 2023-2024 northern hemisphere influenza season

(5) Organization: National Institute for Medical Research

Access: <https://www.crick.ac.uk/partnerships/worldwide-influenza-centre/annual-and-interim-reports>

Detail:

Interim Report February 2024

Interim Report September 2023

Interim Report February 2023

Interim Report September 2022

Interim Report February 2022

Interim Report September 2021

Interim Report February 2021

Interim Report September 2020

Interim Report February 2020

Interim Report September 2019

Interim Report February 2019

Interim Report September 2018

Interim Report February 2018

Interim Report September 2017

Interim Report February 2017

Interim Report September 2016

Interim Report February 2016

Interim Report September 2015

Interim Report February 2015

Interim Report September 2014

Interim Report February 2014

(6) Published papers:

Liu F, Levine M Z. Heterologous Antibody Responses Conferred by A (H3N2) Variant and Seasonal Influenza Vaccination Against Newly Emerged 2016–2018 A (H3N2) Variant Viruses in Healthy Persons [J]. Clinical Infectious Diseases, 2020, 71(12): 3061-3070.

**Supplementary Figures**

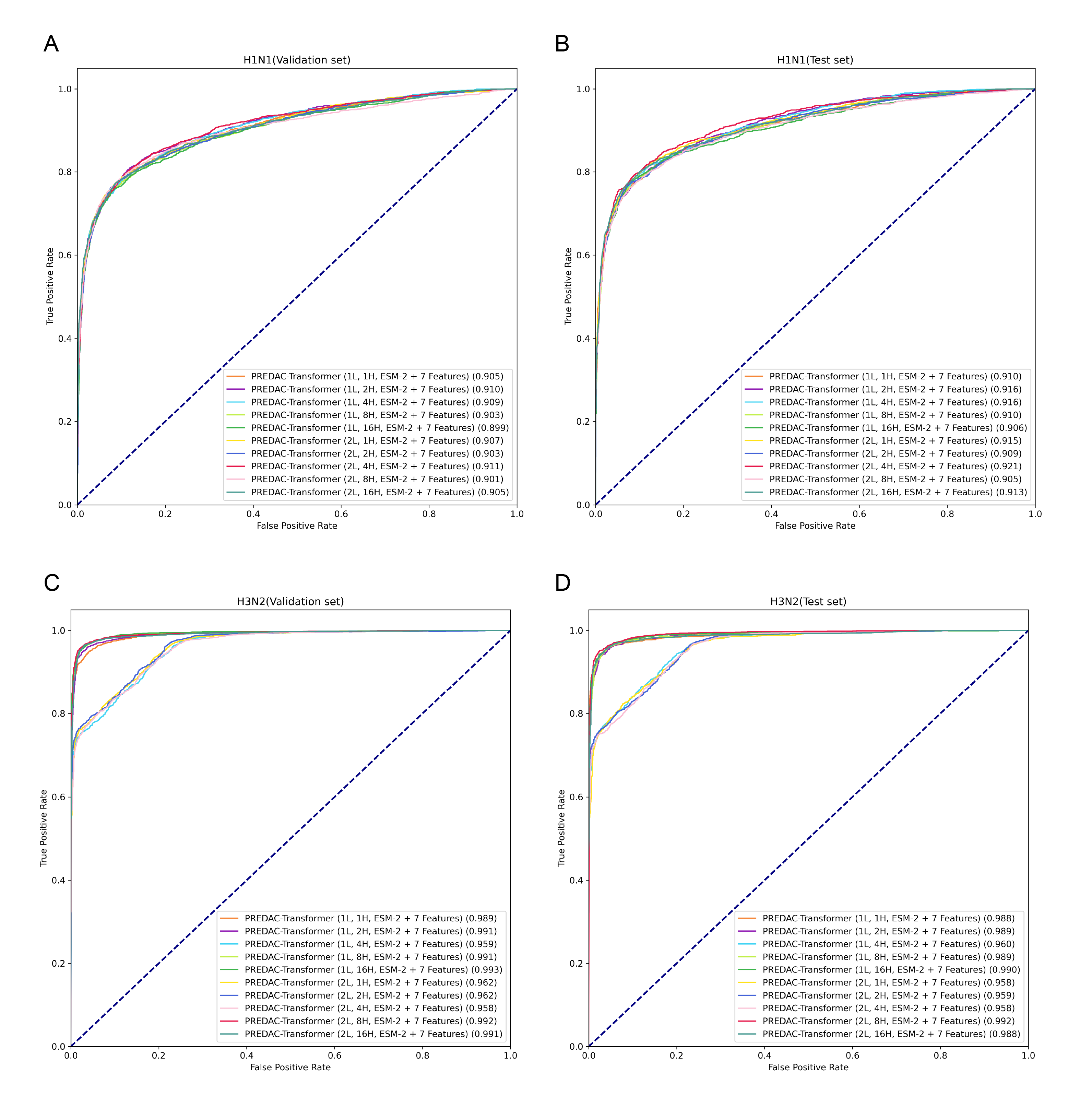

**Supplementary Figure S1.** **Performance of different PREDAC-Transformer model architectures.**

Receiver operating characteristic (ROC) curves evaluating the performance of 10 different Transformer encoder architectures on the task of predicting antigenic relationships for seasonal influenza A viruses. All models were tested using the optimal hybrid feature encoding (ESM-2 + 7-features). (A) ROC curves for influenza A/H1N1 virus pairs on the validation set. (B) ROC curves for influenza A/H1N1 virus pairs on the test set. (C) ROC curves for influenza A/H3N2 virus pairs on the validation set. (D) ROC curves for influenza A/H3N2 virus pairs on the test set. In the legend, 'L' denotes the number of encoder layers and 'H' denotes the number of attention heads. The Area Under the Curve (AUC) value is provided for each architecture to quantify its overall performance.

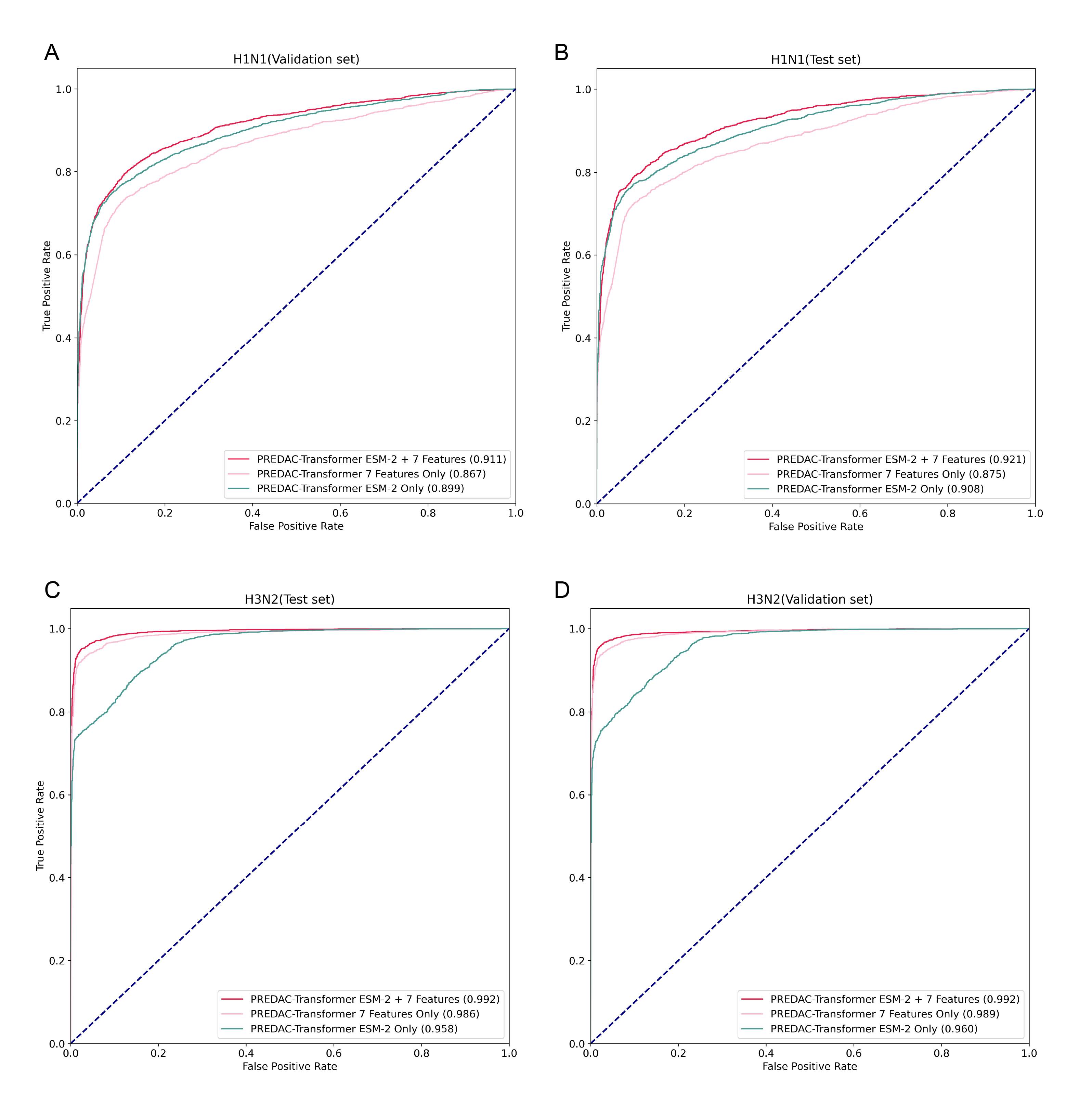

**Supplementary Figure S2. Comparative analysis of feature encoding schemes for PREDAC-Transformer.**

Receiver operating characteristic (ROC) curves from an ablation study evaluating the performance of three distinct feature encoding schemes. This analysis was conducted to determine the contribution of each feature component. The schemes compared are: 7-features only, ESM-2 embeddings only, and the hybrid ESM-2 + 7-features scheme. To provide a consistent basis for comparison, all schemes were evaluated using the pre-determined optimal model architecture for each respective virus subtype: 2 encoder layers with 4 attention heads for H1N1, and 2 encoder layers with 8 attention heads for H3N2. (A) ROC curves for the three schemes on the influenza A/H1N1 validation set. (B) ROC curves on the A/H1N1 test set. (C) ROC curves on the influenza A/H3N2 validation set. (D) ROC curves on the A/H3N2 test set. The Area Under the Curve (AUC) for each scheme is provided in the legend.

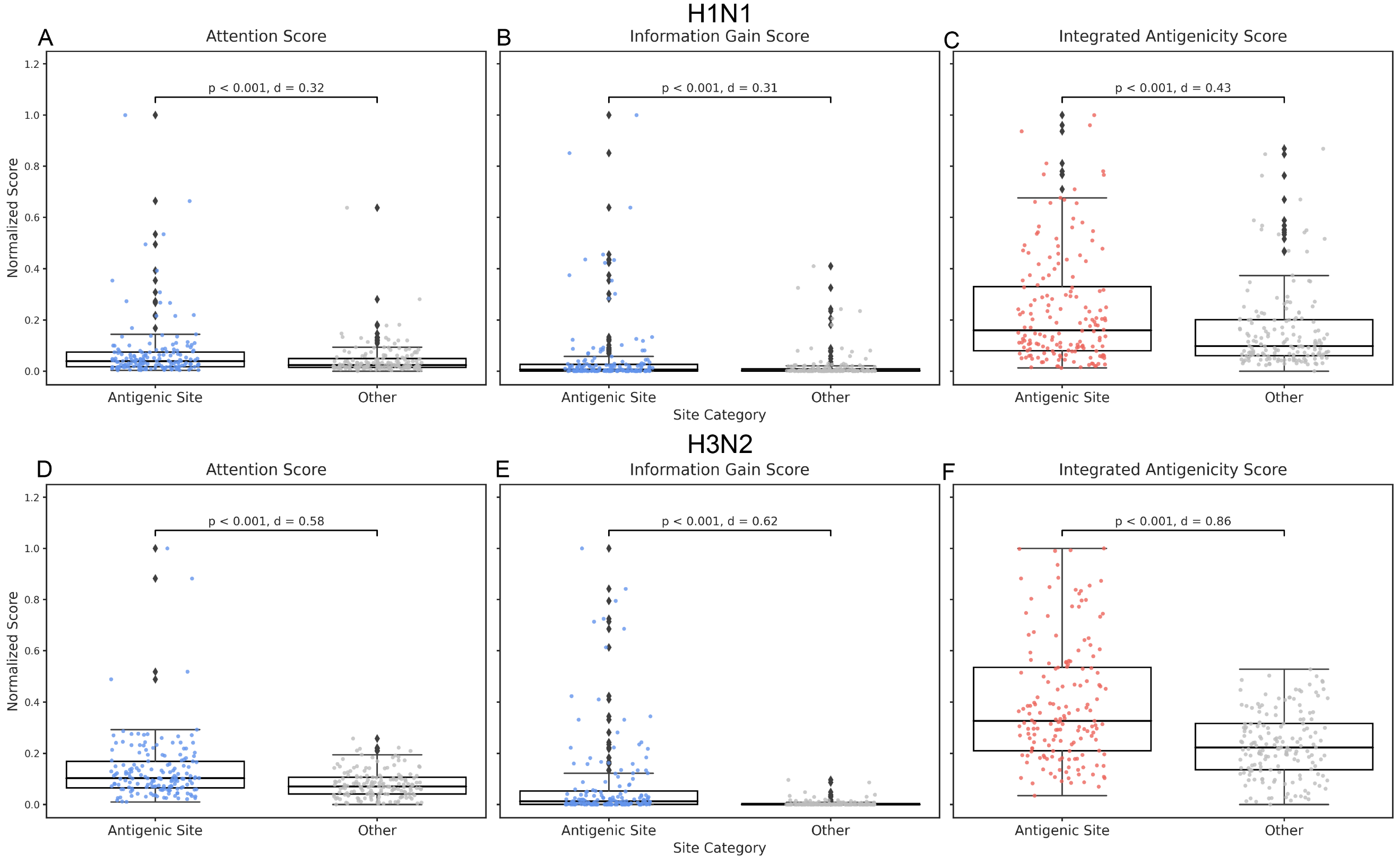

**Supplementary Figure S3. The integrated antigenicity score outperforms individual components in discriminative power.**

This figure presents the results of an ablation study designed to systematically evaluate the individual contributions of the Attention Score and Information Gain to the final integrated antigenicity score, validating the rationale for its integrated design. It compares the ability of these three scores to discriminate between established "Antigenic Sites" and "Other" sites. The top row (A-C) displays the results for the influenza A/H1N1 virus, while the bottom row (D-F) shows the results for the influenza A/H3N2 virus. Each pair of boxplots illustrates the distribution of scores for the two site categories, with statistical significance (p-value from the Mann-Whitney U test) and effect size (Cohen's d) annotated to quantify the difference. Results indicate that integrated antigenicity score outperforms individual scores, demonstrating the highest statistical significance and effect size.

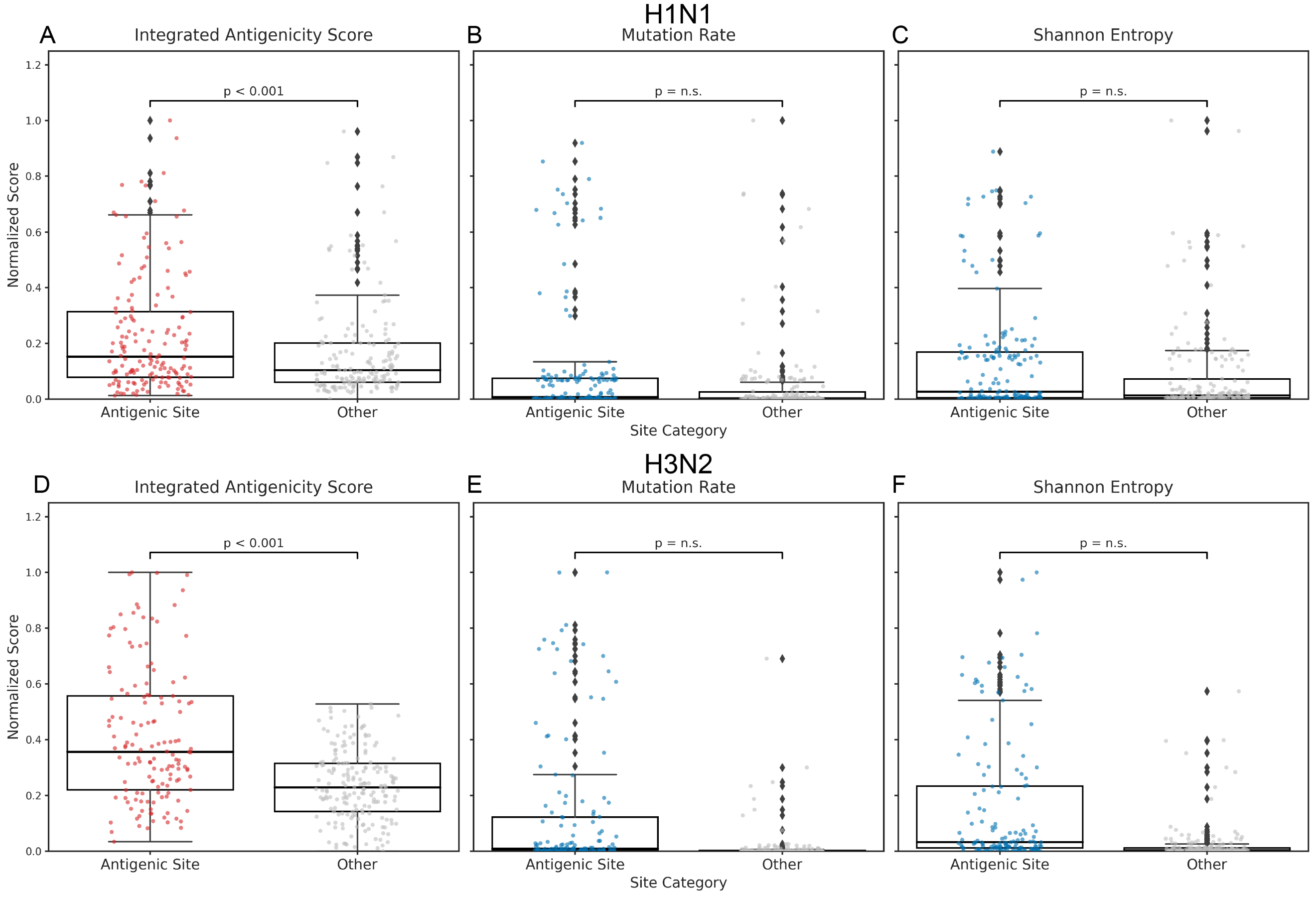

**Supplementary Figure S4. Comparison of the integrated antigenicity score with mutation rate and Shannon entropy**

This figure compares the performance of the integrated antigenicity score with mutation rate and Shannon entropy in discriminating between "Antigenic Sites" and "Other Sites." The top row (A-C) displays results for the influenza A/H1N1 virus, while the bottom row (D-F) shows results for the influenza A/H3N2 virus. Each pair of boxplots illustrates the distribution of scores for the two site categories, with statistical significance (p-value from the Mann-Whitney U test) and effect size (Cohen's d) annotated to quantify the difference.

Results indicate that the integrated antigenicity score outperforms mutation rate and Shannon entropy, demonstrating the highest statistical significance and effect size in distinguishing between the two site categories.

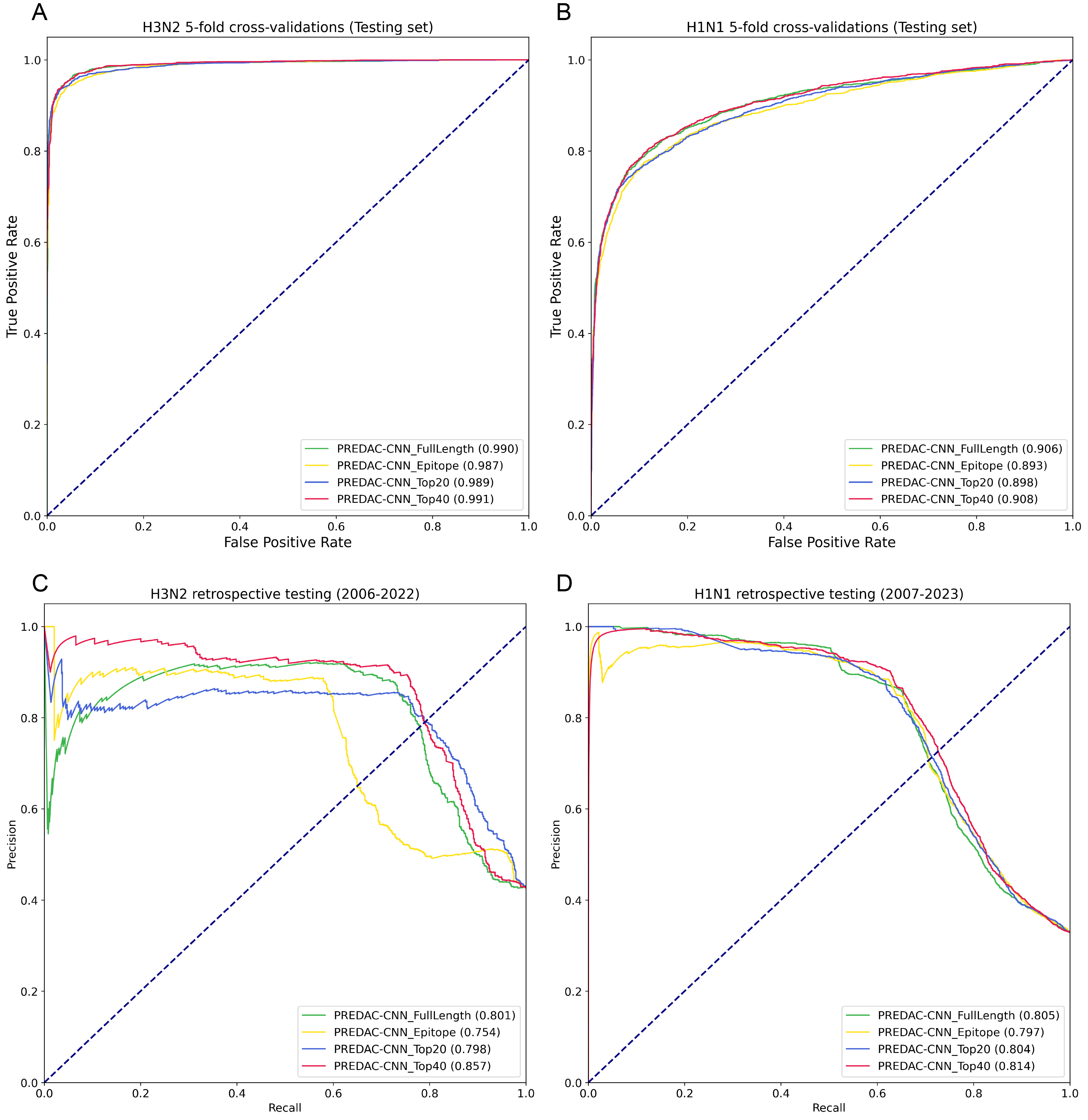

**Supplementary Figure S5. Focusing model attention on global key sites significantly enhances antigenic prediction performance.**

To test whether focusing on key antigenic determinants improves predictive performance, we modified the PREDAC-CNN model by increasing its attention on specific site sets. This figure compares the performance of this weighted model when using four different input feature sets: the full-length HA1 sequence, traditional antigenic epitopes, the top 20 integrated antigenicity score -ranked sites, and the top 40 integrated antigenicity score -ranked sites, which correspond to our definition of global key sites.

The figure displays (A) Receiver Operating Characteristic (ROC) curves for 5-fold cross-validations on influenza A/H3N2 and (B) A/H1N1 viruses. It also shows (C) Precision-Recall (PR) curves for retrospective testing on influenza A/H3N2 and (D) A/H1N1 viruses. The results consistently show that focusing the model on the top 40 global key sites yields superior performance compared to using the full sequence or traditional epitopes, validating global key sites as a compact and potent feature set for enhancing model accuracy.

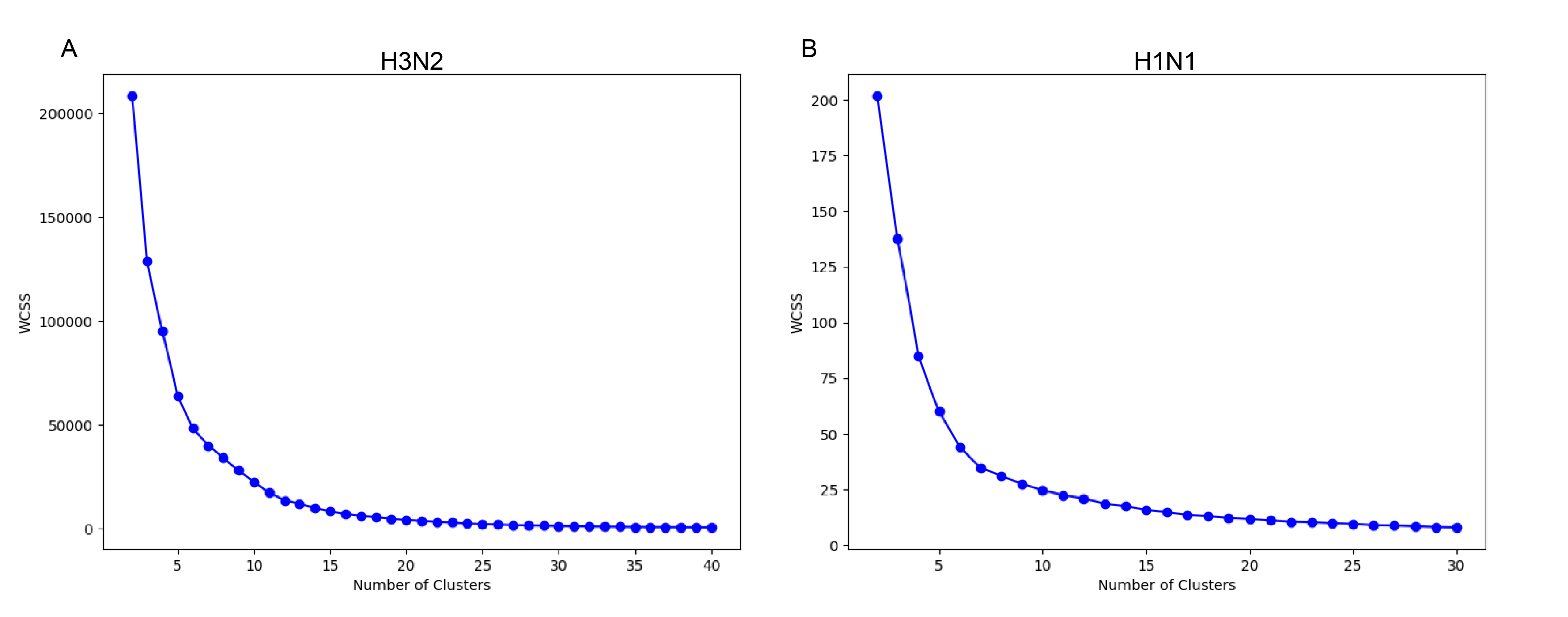

**Supplementary Figure S6. Determination of the optimal number of antigenic clusters using the elbow method**

The figure shows the application of the elbow method to determine the optimal number of clusters (K) for K-means clustering of antigenic data for (A) influenza A/H3N2 and (B) A/H1N1 viruses. For the complete datasets, the elbow method was used to identify the optimal cluster numbers. Additionally, for the sparser, more clustered pre-2009 data, a secondary clustering was performed, with the elbow method indicating the optimal number of clusters.

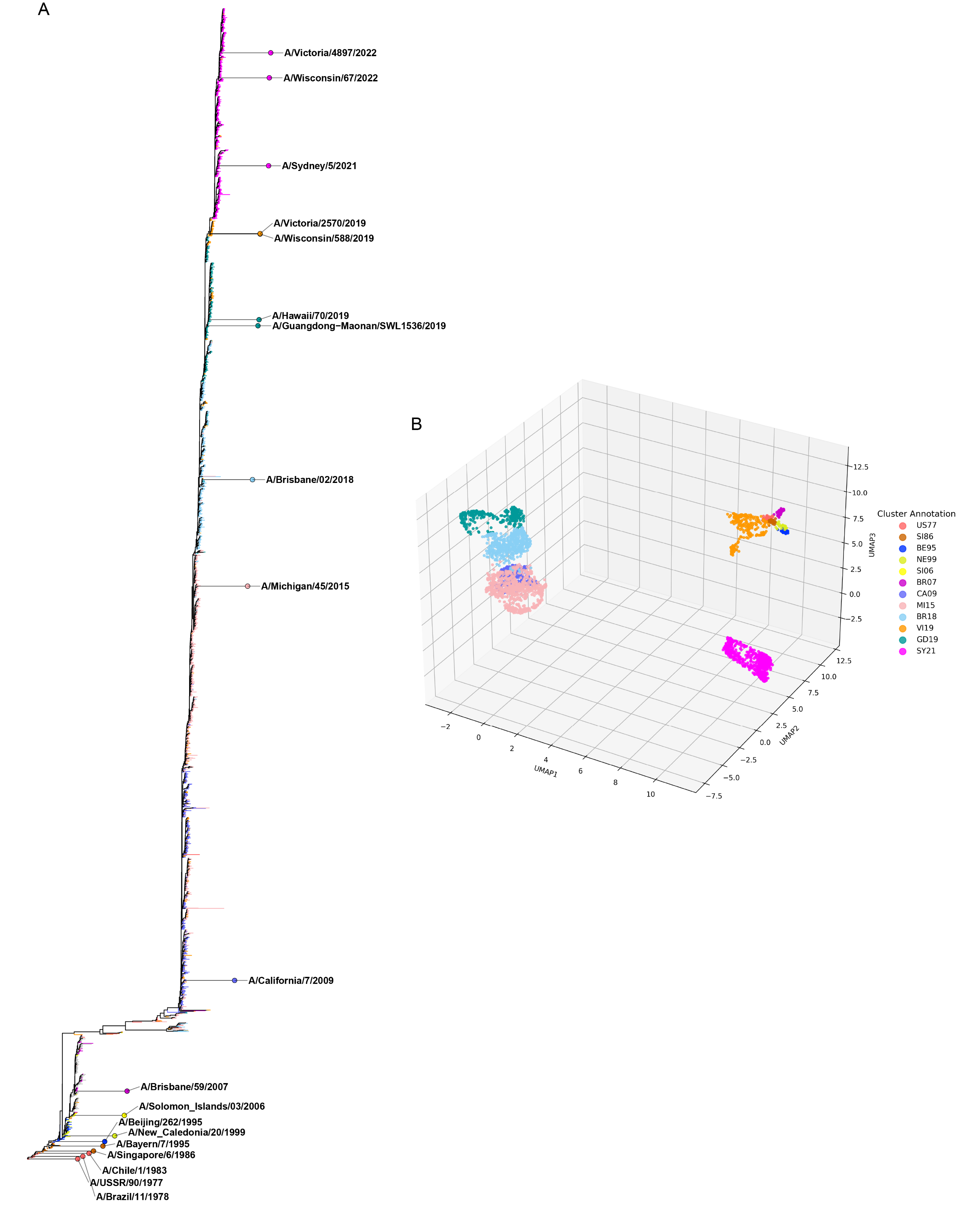

**Supplementary Figure S7. Genetic and antigenic evolution of the influenza A/H1N1 virus.**

(A) The genetic evolution (phylogenetic tree) of the virus, with WHO-recommended vaccine strains denoted by five-pointed stars. (B) The distribution of the corresponding antigenic clusters in a three-dimensional UMAP (Uniform Manifold Approximation and Projection) space. Each antigenic cluster is named using an abbreviation derived from the earliest WHO-recommended vaccine strain it contains.

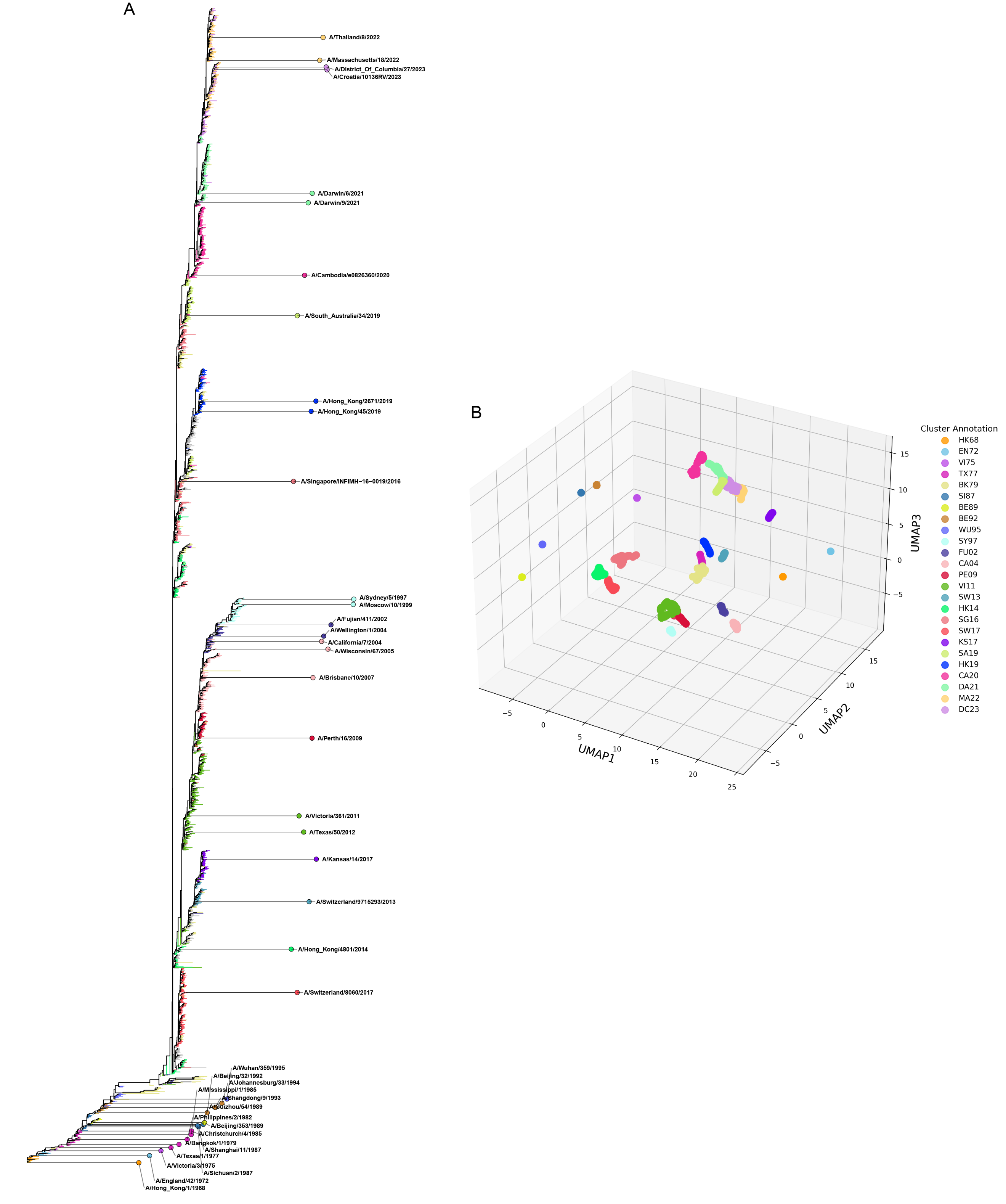

**Supplementary Figure S8. Genetic and antigenic evolution of the influenza A/H3N2 virus.**

(A) The genetic evolution (phylogenetic tree) of the virus, with WHO-recommended vaccine strains denoted by five-pointed stars. (B) The distribution of the corresponding antigenic clusters in a three-dimensional UMAP (Uniform Manifold Approximation and Projection) space. Each antigenic cluster is named using an abbreviation derived from the earliest WHO-recommended vaccine strain it contains.

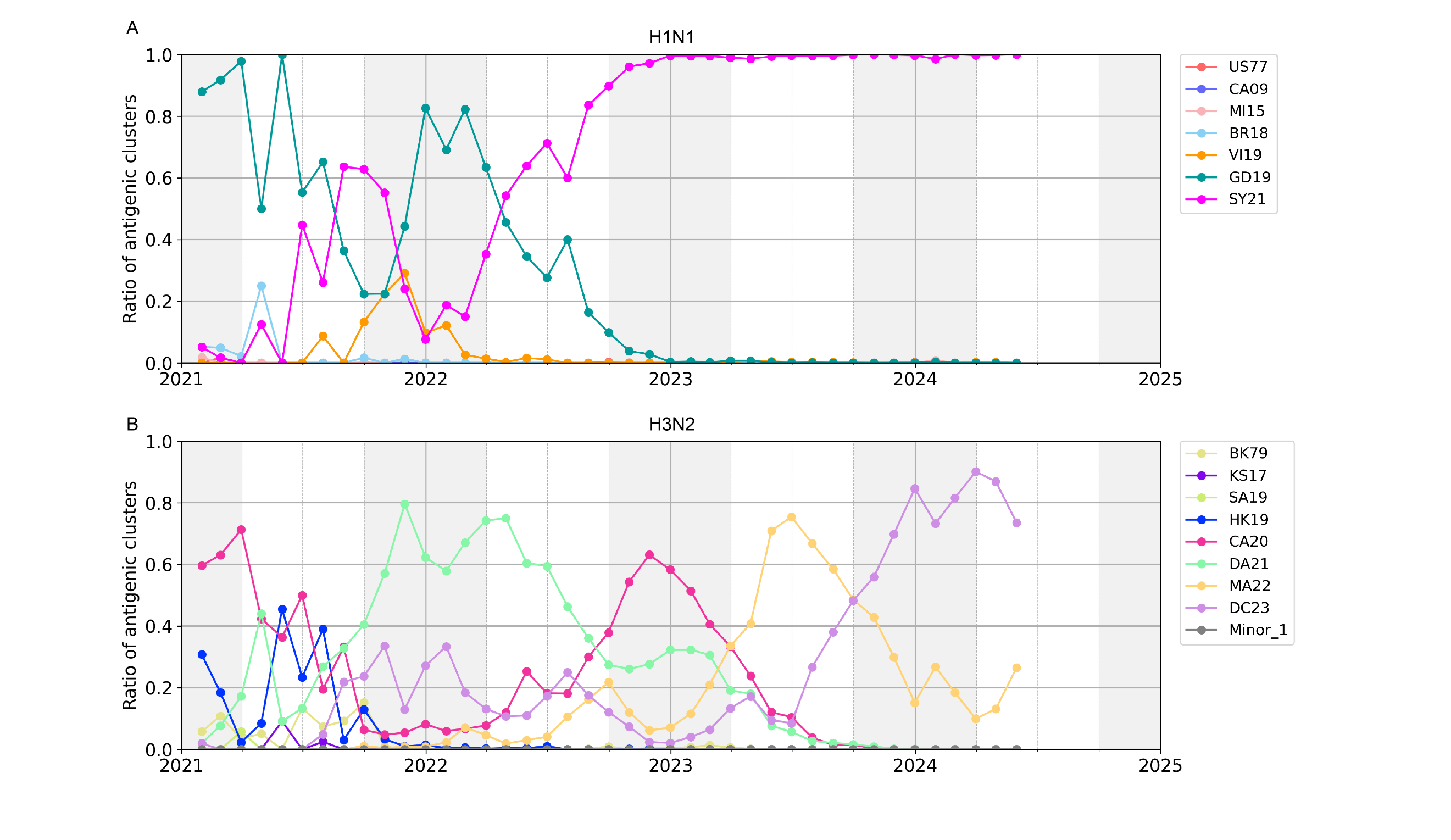

**Supplementary Figure S9. The antigenic clustering and dynamics of influenza A/H1N1 and A/H3N2 viruses (2021–2024).**

This figure illustrates the monthly dynamics of circulating antigenic clusters for (A) influenza A/H1N1 and (B) A/H3N2, based on globally monitored strains from 2021 to 2024. The x-axis represents the year, and the y-axis indicates the proportion of each antigenic cluster. Different antigenic clusters are represented by distinct colored lines, and the grey shaded areas indicate the winter season in the Northern Hemisphere.

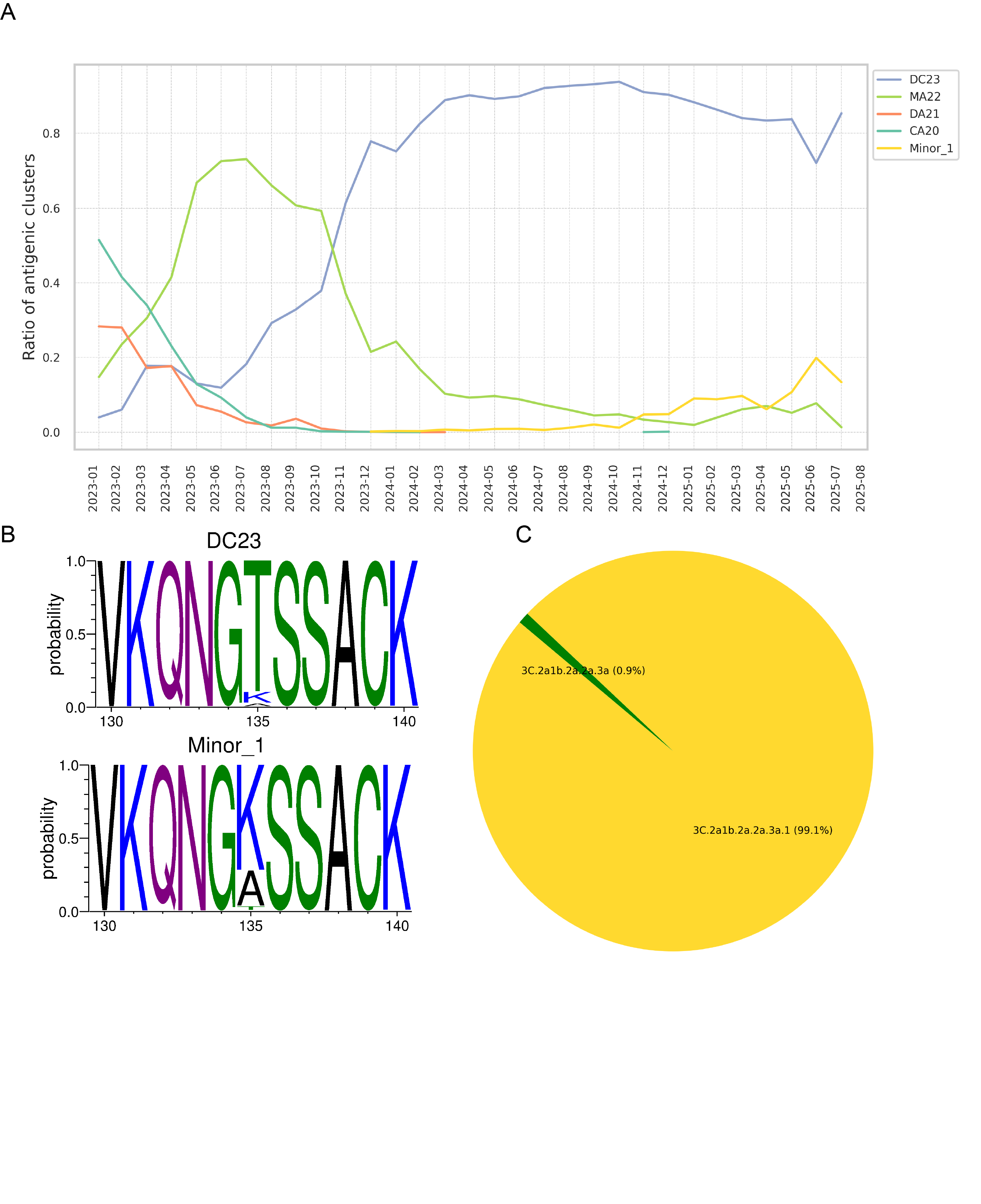

**Supplementary Figure S10. The antigenic clustering of influenza A(H3N2) viruses and analysis of the Minor antigenic cluster.**

(A) Monthly dynamics of antigenic clusters in the periods of 2023-2025. The colored lines refer to the ratio of antigenic clusters, while the gray lines refer to the minor clusters. (B) Analysis of amino acid variation in the DC23 and Minor_1 sequences. The figure highlights the variation at position 135, along with the upstream and downstream 5 amino acid positions. Letters represent amino acids, with heights proportional to their occurrence probability at each position. (C) The clade composition of viruses isolated in 2025 in the Minor_1.

**Supplementary Tables**

**Supplementary Table S1.** The distributions of antigenically distinct strain pairs and antigenically similar strain pairs for each year for influenza A/H3N2 viruses

| Year | Antigenically similar | Antigenically distinct | Year | Antigenically similar | Antigenically distinct |
| --- | --- | --- | --- | --- | --- |
| 1968 | 3 | 0 | 1996 | 203 | 307 |
| 1969 | 7 | 2 | 1997 | 129 | 184 |
| 1970 | 11 | 0 | 1998 | 34 | 41 |
| 1971 | 35 | 4 | 1999 | 66 | 21 |
| 1972 | 12 | 40 | 2000 | 13 | 4 |
| 1973 | 23 | 46 | 2001 | 75 | 4 |
| 1974 | 45 | 26 | 2002 | 44 | 12 |
| 1975 | 22 | 28 | 2003 | 64 | 29 |
| 1976 | 8 | 44 | 2004 | 105 | 25 |
| 1977 | 24 | 25 | 2005 | 115 | 61 |
| 1979 | 0 | 14 | 2006 | 80 | 17 |
| 1980 | 3 | 5 | 2007 | 165 | 20 |
| 1981 | 6 | 0 | 2008 | 44 | 11 |
| 1982 | 18 | 27 | 2009 | 42 | 136 |
| 1983 | 6 | 0 | 2010 | 62 | 80 |
| 1984 | 6 | 19 | 2011 | 56 | 11 |
| 1985 | 59 | 10 | 2012 | 47 | 0 |
| 1986 | 17 | 0 | 2013 | 79 | 37 |
| 1987 | 16 | 81 | 2014 | 59 | 39 |
| 1988 | 34 | 28 | 2015 | 14 | 17 |
| 1989 | 136 | 132 | 2016 | 82 | 50 |
| 1990 | 55 | 60 | 2017 | 42 | 61 |
| 1991 | 76 | 151 | 2018 | 12 | 42 |
| 1992 | 370 | 567 | 2019 | 16 | 47 |
| 1993 | 617 | 1458 | 2020 | 52 | 43 |
| 1994 | 254 | 322 | 2021 | 46 | 32 |
| 1995 | 268 | 471 | 2022 | 119 | 75 |

**Supplementary Table S2.** The distributions of antigenically distinct strain pairs and antigenically similar strain pairs for each year for influenza A/H1N1 viruses

| Year | Antigenically similar | Antigenically distinct |
| --- | --- | --- |
| 1995 | 0 | 1 |
| 1998 | 0 | 1 |
| 2000 | 0 | 1 |
| 2002 | 1 | 0 |
| 2003 | 0 | 1 |
| 2004 | 1 | 1 |
| 2006 | 0 | 1 |
| 2007 | 3 | 10 |
| 2008 | 4 | 12 |
| 2009 | 117 | 66 |
| 2010 | 139 | 104 |
| 2011 | 125 | 103 |
| 2012 | 216 | 110 |
| 2013 | 400 | 180 |
| 2014 | 236 | 95 |
| 2015 | 467 | 247 |
| 2016 | 454 | 245 |
| 2017 | 537 | 100 |
| 2018 | 1547 | 362 |
| 2019 | 1891 | 803 |
| 2020 | 511 | 380 |
| 2021 | 150 | 400 |
| 2022 | 230 | 343 |
| 2023 | 373 | 67 |
| 2024 | 3 | 0 |

**Supplementary Table S3.** Performance comparison of PREDAC-Transformer and its six competitors in 5-fold Cross-Validation of influenza A/H3N2 viruses.

| Model | Accuracy | F1score | Precision | Recall | Specificity | AUC | AUPR |
| --- | --- | --- | --- | --- | --- | --- | --- |
| Transformer | 0.967 | 0.970 | 0.976 | 0.964 | 0.970 | 0.992 | 0.994 |
| PREDAC-CNN | 0.972 | 0.975 | 0.980 | 0.970 | 0.975 | 0.992 | 0.994 |
| IAV-CNN | 0.839 | 0.851 | 0.877 | 0.827 | 0.855 | 0.916 | 0.940 |
| PREDAC-H3 | 0.751 | 0.806 | 0.710 | 0.932 | 0.527 | 0.749 | 0.718 |
| PREDAV-FluA | 0.790 | 0.805 | 0.829 | 0.783 | 0.800 | 0.863 | 0.905 |
| Lee | 0.554 | 0.713 | 0.554 | 1.000 | 0.001 | 0.833 | 0.838 |
| Lees | 0.802 | 0.822 | 0.820 | 0.823 | 0.776 | 0.893 | 0.912 |

**Supplementary Table S4.** Performance comparison of PREDAC-Transformer and its six competitors on independent test sets of influenza A/H3N2 viruses.

| Model | Accuracy | F1score | Precision | Recall | Specificity | AUC | AUPR |
| --- | --- | --- | --- | --- | --- | --- | --- |
| Transformer | 0.960 | 0.964 | 0.967 | 0.961 | 0.959 | 0.992 | 0.993 |
| PREDAC-CNN | 0.956 | 0.960 | 0.958 | 0.963 | 0.947 | 0.990 | 0.992 |
| IAV-CNN | 0.843 | 0.854 | 0.879 | 0.831 | 0.857 | 0.922 | 0.943 |
| PREDAC-H1 | 0.751 | 0.806 | 0.710 | 0.932 | 0.527 | 0.749 | 0.718 |
| PREDAV-FluA | 0.790 | 0.805 | 0.829 | 0.783 | 0.800 | 0.863 | 0.905 |
| Lee | 0.554 | 0.713 | 0.554 | 1.000 | 0.001 | 0.833 | 0.838 |
| Lees | 0.802 | 0.822 | 0.820 | 0.823 | 0.776 | 0.893 | 0.912 |

**Supplementary Table S5.** Performance comparison of PREDAC-Transformer and its six competitors in 5-fold Cross-Validation of influenza A/H1N1 viruses.

| Model | Accuracy | F1score | Precision | Recall | Specificity | AUC | AUPR |
| --- | --- | --- | --- | --- | --- | --- | --- |
| Transformer | 0.868 | 0.787 | 0.839 | 0.740 | 0.930 | 0.911 | 0.875 |
| PREDAC-CNN | 0.868 | 0.793 | 0.820 | 0.767 | 0.917 | 0.895 | 0.865 |
| IAV-CNN | 0.820 | 0.659 | 0.879 | 0.527 | 0.964 | 0.830 | 0.779 |
| PREDAC-H1 | 0.656 | 0.570 | 0.484 | 0.692 | 0.638 | 0.694 | 0.464 |
| PREDAV-FluA | 0.710 | 0.266 | 0.806 | 0.159 | 0.981 | 0.775 | 0.616 |
| Lee | 0.330 | 0.496 | 0.330 | 1.000 | 0.001 | 0.704 | 0.475 |
| Lees | 0.725 | 0.496 | 0.627 | 0.410 | 0.880 | 0.760 | 0.614 |

**Supplementary Table S6.** Performance comparison of PREDAC-Transformer and its six competitors on independent test sets of influenza A/H1N1 viruses.

| Model | Accuracy | F1score | Precision | Recall | Specificity | AUC | AUPR |
| --- | --- | --- | --- | --- | --- | --- | --- |
| Transformer | 0.875 | 0.803 | 0.832 | 0.776 | 0.923 | 0.921 | 0.889 |
| PREDAC-CNN | 0.862 | 0.786 | 0.800 | 0.772 | 0.906 | 0.906 | 0.875 |
| IAV-CNN | 0.819 | 0.665 | 0.847 | 0.547 | 0.952 | 0.829 | 0.767 |
| PREDAC-H1 | 0.656 | 0.570 | 0.484 | 0.692 | 0.638 | 0.694 | 0.464 |
| PREDAV-FluA | 0.710 | 0.266 | 0.806 | 0.159 | 0.981 | 0.775 | 0.616 |
| Lee | 0.330 | 0.496 | 0.330 | 1.000 | 0.001 | 0.704 | 0.475 |
| Lees | 0.725 | 0.496 | 0.627 | 0.410 | 0.880 | 0.760 | 0.614 |

**Supplementary Table S7.** AUPRC (Area Under Precision-Recall Receiver Operating Characteristic) values for PREDAC-Transformer and its five competitors on 17 testing subsets of influenza A/H3N2 viruses.

| Year | AUPRC | | | | | | |
| --- | --- | --- | --- | --- | --- | --- | --- |
|  | Transformer | PREDAC-CNN | IAV-CNN | PREDAC-H3 | PREDAV-FluA | Lee | Lees |
| 2006 | 0.345 | **0.452** | 0.249 | 0.193 | 0.305 | 0.268 | 0.236 |
| 2007 | 0.906 | **0.917** | 0.632 | 0.157 | 0.755 | 0.427 | 0.610 |
| 2008 | 0.471 | 0.470 | 0.605 | 0.252 | 0.452 | **0.634** | 0.625 |
| 2009 | **0.997** | 0.996 | 0.981 | 0.981 | 0.997 | 0.977 | 0.995 |
| 2010 | **0.952** | 0.938 | 0.823 | 0.710 | 0.848 | 0.805 | 0.851 |
| 2011 | **0.955** | 0.795 | 0.175 | 0.227 | 0.644 | 0.184 | 0.462 |
| 2012 | NA | NA | NA | NA | NA | NA | NA |
| 2013 | 0.392 | 0.423 | 0.428 | 0.420 | 0.367 | 0.442 | **0.512** |
| 2014 | 0.646 | 0.577 | 0.628 | 0.543 | **0.751** | 0.667 | 0.566 |
| 2015 | 0.978 | **1.000** | 1.000 | 0.773 | 0.900 | 1.000 | 1.000 |
| 2016 | **0.759** | 0.612 | 0.624 | 0.479 | 0.683 | 0.579 | 0.621 |
| 2017 | 0.845 | 0.877 | 0.831 | 0.677 | 0.761 | 0.850 | **0.888** |
| 2018 | 0.972 | 0.878 | 0.962 | 0.824 | 0.966 | 0.969 | **0.991** |
| 2019 | 0.922 | 0.932 | **0.955** | 0.807 | 0.939 | 0.946 | 0.928 |
| 2020 | 0.549 | **0.618** | 0.555 | 0.450 | 0.516 | 0.572 | 0.539 |
| 2021 | 0.865 | 0.934 | 0.988 | 0.933 | **0.990** | 0.986 | 0.985 |
| 2022 | **0.877** | 0.801 | 0.864 | 0.597 | 0.873 | 0.816 | 0.868 |

**Supplementary Table S8.** AUPRC (Area Under Precision-Recall Receiver Operating Characteristic) values for PREDAC-Transformer and its five competitors on 17 testing subsets of influenza A/H1N1 viruses.

| Year | AUPRC | | | | | | |
| --- | --- | --- | --- | --- | --- | --- | --- |
|  | Transformer | PREDAC-CNN | IAV-CNN | PREDAC-H3 | PREDAV-FluA | Lee | Lees |
| 2007 | 0.925 | 0.708 | 0.900 | **0.967** | 0.783 | 0.904 | 0.885 |
| 2008 | 0.626 | 0.694 | 0.764 | 0.750 | 0.664 | 0.792 | **0.833** |
| 2009 | 0.440 | 0.336 | 0.517 | 0.478 | 0.443 | 0.535 | **0.629** |
| 2010 | 0.685 | 0.662 | 0.539 | 0.468 | 0.508 | 0.525 | **0.708** |
| 2011 | 0.575 | **0.809** | 0.491 | 0.460 | 0.537 | 0.485 | 0.706 |
| 2012 | **0.850** | 0.850 | 0.420 | 0.380 | 0.435 | 0.386 | 0.769 |
| 2013 | 0.870 | **0.874** | 0.462 | 0.411 | 0.459 | 0.397 | 0.835 |
| 2014 | **0.846** | 0.766 | 0.716 | 0.446 | 0.414 | 0.402 | 0.738 |
| 2015 | 0.779 | **0.798** | 0.789 | 0.442 | 0.520 | 0.421 | 0.764 |
| 2016 | **0.825** | 0.809 | 0.756 | 0.442 | 0.552 | 0.427 | 0.719 |
| 2017 | **0.560** | 0.404 | 0.257 | 0.265 | 0.214 | 0.209 | 0.244 |
| 2018 | **0.568** | 0.544 | 0.379 | 0.307 | 0.342 | 0.313 | 0.300 |
| 2019 | **0.755** | 0.732 | 0.607 | 0.452 | 0.502 | 0.567 | 0.454 |
| 2020 | **0.917** | 0.906 | 0.684 | 0.498 | 0.632 | 0.598 | 0.419 |
| 2021 | **0.986** | 0.981 | 0.932 | 0.864 | 0.962 | 0.937 | 0.921 |
| 2022 | 0.948 | 0.930 | 0.937 | 0.897 | **0.963** | 0.932 | 0.900 |
| 2023 | **0.602** | 0.499 | 0.572 | 0.499 | 0.571 | 0.532 | 0.482 |
